# Ehrlich occupancy time: Beyond *k*_off_ to a complete residence time framework

**DOI:** 10.64898/2025.12.10.693538

**Authors:** Justin Eilertsen, Santiago Schnell, Sebastian Walcher

## Abstract

Drug-target occupancy time—the cumulative duration a target remains bound—critically influences therapeutic efficacy. While Copeland’s widely-used residence time (1*/k*_off_) emphasizes dissociation kinetics, it neglects association rates, rebinding events, and drug elimination that affect in vivo outcomes. Returning to Paul Ehrlich’s 1913 principle that drugs act only when bound (“Corpora non agunt nisi fixata”), we develop a mathematically rigorous framework defining Ehrlich occupancy time, EOT, as the integral of fractional target occupancy over time. Our approach explicitly incorporates association (*k*_on_) and dissociation (*k*_off_) kinetics, accounts for rebinding, and extends to systems with drug removal. For drug-receptor closed systems at equilibrium, we prove that relative EOT equals the equilibrium occupancy fraction; under ligand-excess conditions this reduces to *b*_0_*/*(*K*_*d*_ +*b*_0_), where *K*_*d*_ is the dissociation constant and *b*_0_ the drug concentration. For induced-fit mechanisms, conformational changes reduce the effective dissociation constant to 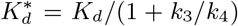 (where *k*_3_ and *k*_4_ are forward and reverse isomerization rates), prolonging occupancy through kinetic trapping. Critically, for drug-receptor systems with firstorder drug elimination at rate *k*_3_, we derive rigorous bounds: *b*_0_*/*[(*b*_0_ + *K*_*d*_)*k*_3_] ≤ EOT_∞_ ≤ *b*_0_*/*(*K*_*d*_ · *k*_3_), where EOT_∞_:= lim_*T*→∞_ EOT(*T*) is the total cumulative occupancy time as *T*→ ∞, revealing that both binding affinity and elimination rate jointly determine occupancy. This explains why high-affinity drugs can fail clinically if eliminated rapidly, and identifies pharmacokinetic optimization opportunities. We prove Copeland’s definition is a special case of Ehrlich occupancy time when rebinding is absent. Our framework provides quantitative tools for optimizing drug design beyond binding affinity and enables improved prediction of in vivo efficacy where pharmacokinetics dominate.

## 1 Introduction

The development of effective therapeutics requires understanding not merely how strongly drugs bind to their targets, but how long they remain bound. Traditional drug discovery has emphasized binding affinity— quantified by the equilibrium dissociation constant *K*_*d*_—as the primary metric for drug-target interactions. However, binding affinity alone often fails to predict therapeutic efficacy [1, 2]. Drugs with moderate affinity may exhibit prolonged effects due to extended binding durations, while high-affinity compounds can dissociate rapidly, yielding short-lived responses [3, 4].

The insight that the *duration* of drug-target binding critically determines drug efficacy traces to Paul Ehrlich’s 1913 principle [5]: “Corpora non agunt nisi fixata” –substances act only when bound. Ehrlich recognized that therapeutic effect depends on the cumulative time a target remains occupied by drug molecules. Building on this foundation, Copeland and colleagues introduced “drug-target residence time” (DTRT), defined as the reciprocal of the dissociation rate constant: DTRT = 1*/k*_off_ [1, 6, 7]. This concept has influenced modern drug design, particularly for kinase inhibitors [8] and antibiotics [9, 10], where prolonged target engagement correlates with enhanced efficacy.

However, Copeland’s DTRT definition has significant limitations. First, it considers only dissociation kinetics, neglecting the association rate (*k*_on_) and thereby ignoring how quickly drugs initially engage targets. Second, because 1*/k*_off_ is the mean duration of a *single* binding event, it does not by itself reflect the additional occupancy generated when dissociated drug molecules reassociate with the same or nearby targets, which can substantially extend cumulative target engagement in vivo [11]. Third, DTRT assumes equilibrium conditions and does not account for drug removal processes (metabolism, excretion) that dominate in physiological settings. Folmer [12] critiqued DTRT as “misleading,” noting that its predictive value often reduces merely to potency and that claims of benefits beyond drug clearance lack robust experimental support.

The vantage point of the present paper, as in the work of Copeland and colleagues, is Ehrlich’s original principle in verbal form. In contrast to Copeland and colleagues, who consider only part of the reactions involved, we proceed to develop in a rigorous manner, the mathematical underpinning of Ehrlich’s principle, which we term **Ehrlich occupancy time (EOT)**—the total cumulative duration that target molecules are occupied by drug, explicitly including rebinding events and drug removal. Our approach defines EOT rigorously through chemical kinetics, incorporating both *k*_on_ and *k*_off_; quantifies EOT in closed systems, showing that relative EOT equals the equilibrium fraction of bound receptors and that *K*_*d*_ is the appropriate metric; derives analytical bounds for EOT in systems with first-order drug removal, revealing how pharmacokinetic parameters constrain therapeutic duration; extends the framework to induced-fit mechanisms; and proves that EOT reduces to Copeland’s DTRT only in the absence of rebinding.

While our framework is general and applicable to diverse drug-target binding scenarios, we demonstrate its utility through three clinically relevant exemplary cases: (i) simple receptor-ligand binding in closed systems, (ii) induced-fit mechanisms with conformational changes, and (iii) systems with first-order drug removal. These examples span the range from idealized biochemical assays to physiologically relevant in vivo conditions.

Our framework addresses key questions: How do association and dissociation kinetics jointly determine occupancy time? When does drug removal limit therapeutic duration? How do conformational changes in the target affect residence time? The answers provide quantitative guidance for drug design and more accurate prediction of in vivo efficacy.

For context, we note related literature: Copeland’s residence time applies well to cases like kinase inhibitors [8], but in complex systems, rebinding and pharmacokinetics necessitate broader views [11]. Our framework integrates kinetics holistically. Detailed derivations are in appendices, with main text focusing on implications.

This paper is organized as follows. Section 2 develops the mathematical definition of EOT. Section 3 applies the framework to closed systems (receptor-ligand binding, induced fit). Section 4 analyzes systems with drug removal, deriving computable bounds. Section 5 discusses implications for drug discovery. Our framework integrates kinetics holistically, with detailed derivations provided in the appendices and the main text focusing on implications.

## 2 Mathematical Definition of Ehrlich Occupancy Time

### 2.1 Conceptual Foundation and Discrete-Time Formulation

Consider a population of target receptors interacting with drug molecules over an observation period [0, *T*]. At any time *t*, each receptor is either free (unoccupied) or bound (occupied by a drug molecule). The **Ehrlich occupancy time (EOT)** is the average cumulative duration that receptors spend in the bound state during [0, *T*].

The key difference between EOT and DTRT lies in the mathematical formulation of Ehrlich’s concept: on the one hand, EOT provides a complete mathematical framework; on the other hand, the mathematical formulation of DTRT is certainly incomplete, because it does not include any rebinding and—taken literally— would count only the first binding–unbinding event.

For illustration, consider Figure 1C–D. At this stage no assumption on reaction mechanisms is necessary; one uses only that, at any instant, every receptor is either occupied or free. Each row is a schematic occupancy record for a single receptor (among an ensemble of identical ones), drawn only to illustrate the counting convention: EOT sums all occupied intervals, whereas Copeland residence time counts the first alone. In record 1, for instance, the occupied intervals fall in three separate periods, all of which contribute to EOT, while Copeland residence time would count only the first. The formal construction in Section 2.1.1 below likewise rests on no reaction mechanism.

**Figure 1.**
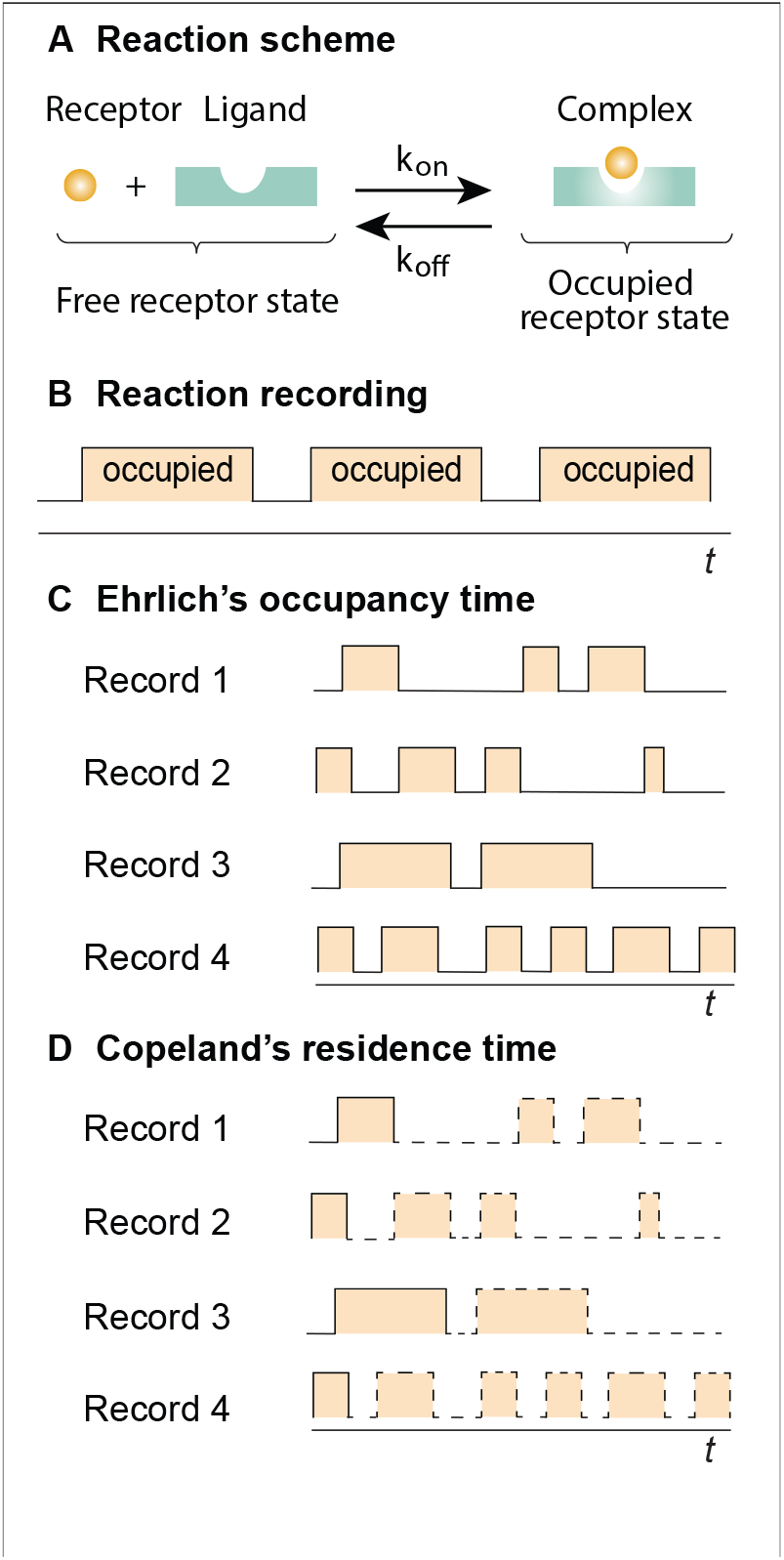
Conceptual comparison of Ehrlich occupancy time (EOT) and Copeland residence time. **(A)** Reversible drug-target binding with association rate *k*_on_ and dissociation rate *k*_off_. **(B)** A generic occupancy recording in which shaded intervals indicate the occupied state. **(C)** EOT bookkeeping: all shaded intervals in an illustrative record are counted, including re-occupancy after prior dissociation. **(D)** Copeland residence-time bookkeeping: only the first occupied interval is counted (solid shading); subsequent occupied intervals are shown with dashed outlines and are excluded. **Key distinction:** EOT measures cumulative target occupancy, whereas Copeland residence time—taken literally—measures a single binding-event duration. The rows labelled Record 1–4 are illustrative single-receptor occupancy records that fix the counting convention. The records are not based on any specific reaction mechanism.

At the single-receptor level, occupancy is a two-state stochastic process. The deterministic, populationlevel fractional occupancy *f* (*t*), defined in equation (4) below and analysed in the remainder of the paper, is the ensemble average of this single-receptor process and, equivalently, the large-*N* limit of that occupancy fraction.

#### 2.1.1 Formal Construction

The following finite-*N* construction and derivation is independent of reaction mechanisms. The only requirement is that each receptor is in an occupied, or non-occupied state, at any given time.

We formalize these concepts mathematically. Consider a finite set of receptors

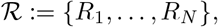

where each receptor *R*_*i*_ can be in one of two states at any time *t* ≥ 0: occupied (bound by a ligand) or free (unoccupied). We observe the system over the interval [0, *T*], which we partition into *M* discrete time points:

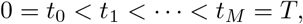

with uniform spacing *t* = *t*_*j*+1_ − *t*_*j*_ = *T/M* for all *j* = 0, 1, …, *M* 1.

The occupancy status of each receptor at each observation time is captured by the indicator function:

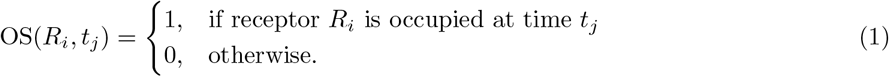

For an individual receptor *R*_*i*_, assuming the occupancy status remains constant within each interval (*t*_*j* − 1_, *t*_*j*_], the total time spent in the occupied state is approximated by:

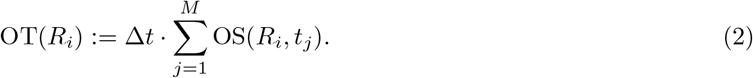

At each time point *t*_*j*_, the number of occupied receptors is

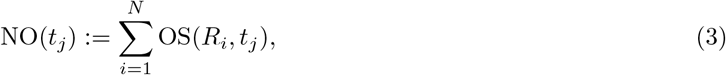

and the fraction of occupied receptors is

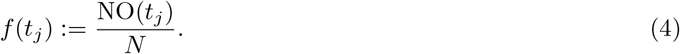

The **Ehrlich occupancy time** is defined as the average occupancy time across all receptors:

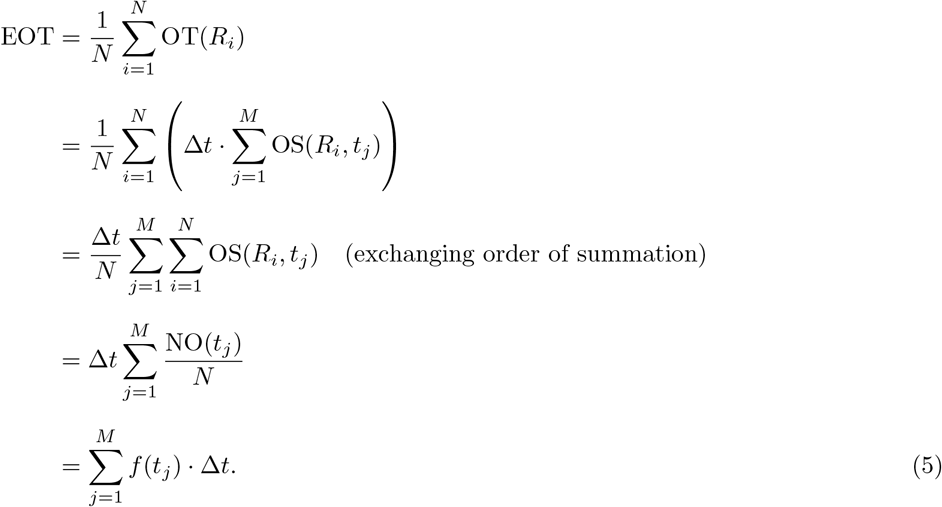

Equation (5) expresses the discrete-time EOT as a weighted sum of the fractional occupancy at each observation time. This formulation has an important consequence: *EOT can be computed from the population-level quantity f* (*t*_*j*_) *without tracking individual receptors*. This connects the microscopic definition (averaging over individual receptor trajectories) to the macroscopic observable (fraction of receptors bound at each time).

##### Example 1

(Discrete-Time Illustration). *Consider N* = 3 *receptors observed at M* = 4 *time points over period T* = 4 *time units, so Δt* = 1. *Suppose the occupancy status is recorded as:*

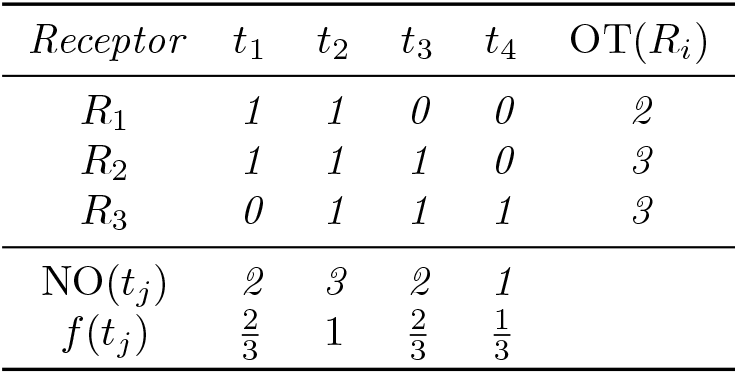

*The EOT can be computed either by averaging individual receptor occupancy times*,

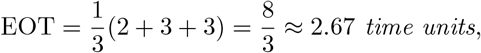

*or equivalently, by summing the fractional occupancies*,

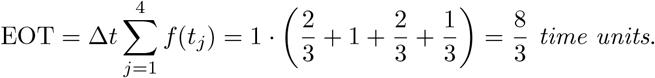

*Both approaches yield the same result, confirming the equivalence established in equation* (5).

##### Remark 1

(Dependence on Δ*t*). *The discrete-time formulation* (5) *depends on the choice of time step Δt. For a fixed observation period T*, *finer temporal resolution (smaller Δt, larger M) generally provides a more accurate approximation of the true cumulative occupancy. In the next subsection, we take the limit t* 0 *to obtain the continuous-time formulation, which eliminates this dependence*.

### 2.2 Continuous-Time Formulation

The discrete-time EOT in equation (5) depends on the choice of partition, and should be viewed as an approximation whose accuracy improves as Δ*t* decreases. To obtain a formulation independent of the observation frequency, we pass to the continuous-time limit.

#### 2.2.1 From Discrete to Continuous Time

Consider the general setting of free receptors (*X*_*i*_), bound complexes (*Y*_*j*_), and other species including ligands (*Z*_*k*_). The concentrations of these species evolve according to a system of ordinary differential equations derived from the reaction kinetics.^1^ The fraction of occupied receptors at any time *t* is given by:

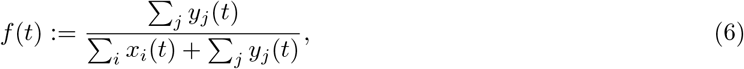

where lowercase letters denote concentrations. In closed systems, the total receptor concentration ∑_*i*_ *x*_*i*_ + ∑_*j*_ *y*_*j*_ is conserved, so the denominator remains constant.

Recall from equation (5) that the discrete-time EOT is given by:

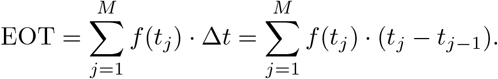

This expression is a *Riemann sum* for the function *f* over the interval [0, *T*], where the time points *t*_0_ *< t*_1_ *<* · · · *< t*_*M*_ form a partition with uniform subinterval length Δ*t* = *T/M*.

A fundamental result in analysis [13] states that if *f*: [0, *T*] → ℝ is continuous, then the Riemann sum converges to the definite integral as the partition is refined:

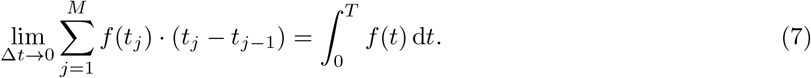

Equivalently, taking *M* ⟶ ∞ while holding *T* fixed (so that Δ*t* = *T/M* → 0) yields the same limit.

When the concentrations *x*_*i*_(*t*) and *y*_*j*_(*t*) are governed by systems of ordinary differential equations with continuous right-hand sides, the solutions are continuous functions of time, and hence *f* (*t*) as defined in (6) is continuous on [0, *T*]. The convergence (7) therefore applies, justifying the following definition.

##### Definition 1

(Ehrlich Occupancy Time). *The* ***Ehrlich occupancy time*** *over the interval* [0, *T*] *is defined as:*

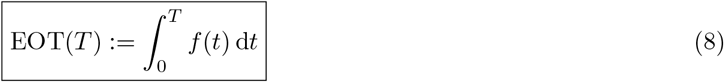

*where f* (*t*) *is the fraction of receptors occupied at time t*.

Equation (8) is our primary definition and serves as the foundation for all subsequent analysis. It has a natural interpretation: EOT(*T*) is the area under the fractional occupancy curve from time 0 to time *T*. The discrete-time formulation (5) provides an approximation to this integral, with accuracy improving as Δ *t* → 0.

#### 2.2.2 Connection to Experimental Measurements

While Equation (8) is defined theoretically, it connects directly to experimentally measurable quantities. The fraction *f* (*t*) can be determined experimentally using several complementary techniques. Surface plasmon resonance (SPR) provides real-time measurement of bound complex concentration, yielding *f* (*t*) directly when the response is normalized by total binding capacity. Radioligand binding assays measure bound radioactivity at multiple time points to construct *f* (*t*) profiles. For live-cell applications, fluorescence-based target engagement assays using FRET or fluorescence polarization can monitor the bound fraction in real time. Cellular thermal shift assays (CETSA) offer an alternative approach, indirectly reporting target occupancy through the stabilization of target proteins against thermal denaturation.

Experimental determination of EOT thus requires measuring *f* (*t*) at multiple time points over the observation period [0, *T*] and numerically integrating. The discrete-time formula (5) corresponds to a left-endpoint Riemann sum; in practice, the trapezoidal rule or Simpson’s rule may provide better accuracy for a given number of measurements. Alternatively, if binding kinetics are well-characterized (known rate constants), one can predict *f* (*t*) computationally and calculate EOT analytically or numerically. Detailed experimental protocols for measuring kinetic parameters and validating EOT predictions are provided in Appendix B.

In Sections 3–4, we evaluate the integral in (8) for specific binding mechanisms to obtain closed-form expressions or computable bounds for EOT.

### 2.3 Relative Ehrlich Occupancy Time

For closed systems approaching equilibrium, the fractional occupancy *f* (*t*) converges to a limiting value *f*_*∞*_ as *t* ⟶ ∞. In this setting, EOT(*T*) grows without bound as *T* increases, making it useful to consider a normalized quantity.

#### Definition 2

(Relative Ehrlich Occupancy Time). *The* ***relative Ehrlich occupancy time*** *over the interval* [0, *T*] *is defined as:*

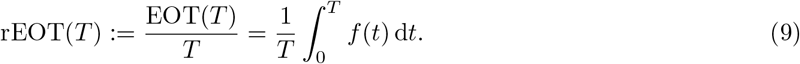

The relative EOT represents the time-averaged fractional occupancy over [0, *T*]. The following proposition establishes that, for systems approaching equilibrium, this average converges to the equilibrium occupancy.

#### Proposition 1

*Let f*: [0, ∞) ⟶ [0, 1] *be a continuous function satisfying f* (*t*) ⟶ *f*_∞_ *as t* → ∞ *for some f*_∞_ ∈ [0, 1]. *Then*

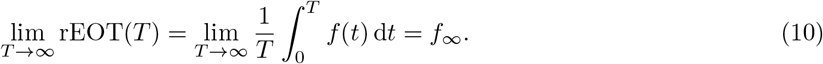

*Proof*. The following proof was suggested by a reviewer; it is more concise than our original one. It uses Lebesgue theory of integration (see for instance Rudin [14]). Via the change of variables *t* = *xT* in the integral defining rEOT(*T*) one gets

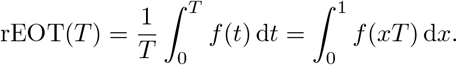

Since *f* (*xT*) ⟶ *f*_∞_ pointwise for every *x* ∈ (0, 1] as *T* → ∞, and |*f* (*xT*)| ≤ 1 provides a dominating integrable function on [0, 1], Lebesgue’s Dominated Convergence Theorem gives 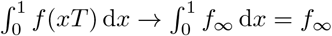.

Proposition 1 provides a direct connection between residence time and target occupancy: *for systems at equilibrium, the relative EOT equals the equilibrium fraction of bound receptors*. This result justifies the use of *f*_*∞*_ as an approximation for rEOT(*T*) when *T* is sufficiently large and the system has equilibrated.

#### Remark 2

(Practical Implications). *Relative EOT is particularly* useful *for comparing drugs in* in vitro *binding assays where the system reaches equilibrium. Since EOT*(*T*) *grows approximately linearly with time in equilibrated systems (EOT*(*T*) ≈ *f*_*∞*_ · *T for large T), the ratio EOT*(*T*)*/T provides a time-independent metric that depends only on the binding kinetics and drug concentration. This approximation is meaningful whenever f*_*∞*_ *>* 0; *in other words, when bound states persevere as t*→∞. *The degenerate case f*_*∞*_ = 0 *can arise in hypothetical scenarios, for instance when a ligand-receptor complex dissociates irreversibly to other species, but these seem of little relevance*.

*In Section 3, we derive explicit expressions for f*_*∞*_ *(and hence rEOT) for several binding mechanisms. For systems with drug removal (Section 4), where equilibrium is never reached and f* (*t*) → 0 *as t* → ∞, *one must instead analyze the limiting value* EOT_*∞*_ = lim_*T→∞*_ EOT(*T*), *which remains finite*.

### 2.4 Relationship to Copeland DTRT

A natural question is how our framework relates to Copeland’s widely-used definition DTRT = 1*/k*_off_ [1]. We now show that Copeland’s metric emerges as a special limiting case of EOT when rebinding is absent.

#### 2.4.1 Derivation of Copeland DTRT as a Limiting Case

We consider a drug–receptor complex that is pre-formed at *t* = 0, with initial concentration *c*(0) = *c*_0_, and study its subsequent decay when reassociation is absent (*k*_1_ = 0). Physically, this corresponds to conditions of infinite dilution or rapid removal of dissociated ligand, so that the probability of a freed drug molecule re-encountering a receptor is negligible (see also [1], Box 1 on page 731). The initial binding event is encoded entirely in the initial condition; the dynamics model only what happens *after* the complex has formed. This situation is modeled by irreversible dissociation:

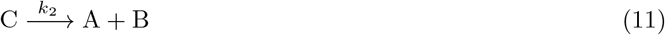

with no reverse (association) reaction, corresponding to *k*_1_ = 0. The complex therefore decays according to first-order kinetics:

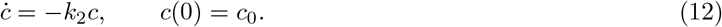

The solution is exponential decay:

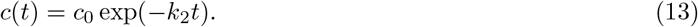

Taking the receptor concentration as *a*_0_ = *c*_0_ (all receptors initially bound), the fractional occupancy is *f* (*t*) = *c*(*t*)*/c*_0_ = exp(−*k*_2_*t*). The total Ehrlich occupancy time is:

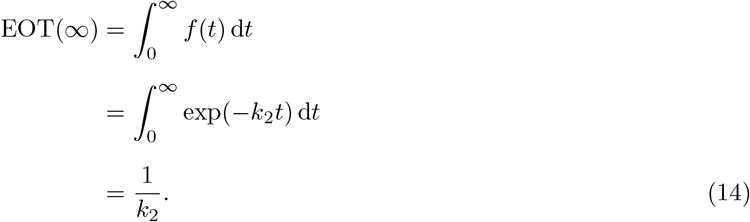

This is precisely Copeland’s definition: DTRT = 1*/k*_off_, where *k*_off_ = *k*_2_ is the dissociation rate constant.

#### 2.4.2 Interpretation and Implications

This derivation has two important implications. First, it establishes that Copeland DTRT is a special case of EOT: when rebinding is absent (*k*_1_ = 0), the Ehrlich occupancy time reduces exactly to the Copeland metric, demonstrating that our framework generalizes, rather than contradicts, the existing definition. Second, the derivation shows that rebinding extends occupancy time. In general binding scenarios (Section 3) where *k*_1_ *>* 0, dissociated drug molecules can reassociate with receptors, contributing additional occupied time. Consequently, EOT exceeds Copeland DTRT, and the difference between EOT and 1*/k*_2_ quantifies the contribution of rebinding to cumulative target occupancy.

Referring to Figure 1, Copeland DTRT corresponds to measuring only the first shaded region for each receptor (panel D), while EOT measures all shaded regions (panel C). The two metrics coincide only when there is at most one shaded region per receptor—that is, when rebinding does not occur.

It is worth making precise in what sense 1*/k*_off_ does, and does not, register rebinding, since this is sometimes a source of confusion. The reciprocal off-rate 1*/k*_off_ is the mean duration of a *single* occupied interval—the mean dwell time per binding event. Operationally it is fixed by the standard dissociation assay, in which an ensemble of pre-formed complexes is allowed to dissociate under sink (non-rebinding) conditions and the bound fraction decays exponentially with rate *k*_off_. The same value is the mean bound-state lifetime in any model in which an occupied target is vacated only by first-order dissociation, since the lifetime of a formed complex is set by *k*_off_ alone and is independent of the surrounding free-ligand concentration; rebinding generates additional occupied intervals without altering the mean duration of each. (For a single receptor held at constant ligand concentration, occupancy is a stationary two-state process and this lifetime is equivalently the long-run time average of the persistence times. In non-closed systems the free ligand is depleted and the process is transient, so the operative statement is this per-interval one rather than a stationary time average.) Cumulative occupancy, by contrast, is the product of the number of occupied intervals and their mean duration, and 1*/k*_off_ reports only the latter factor. Copeland residence time, DTRT = 1*/k*_off_, is therefore a single-event quantity: it is correctly defined and correctly measured, but incomplete as a measure of cumulative occupancy. It does not represent the additional occupancy contributed by re-association, because re-occupancy enters through the *number* of occupied intervals, to which a per-event mean is by construction insensitive. Ehrlich occupancy time captures the full product, and the surplus EOT −1*/k*_off_ is exactly the rebinding contribution (Equation (14) and Section 3). The first-order rate constants in the problem play distinct roles, which are worth separating. Dissociation (*k*_off_, equivalently *k*_diss_) vacates an intact target and sets the per-event occupied lifetime 1*/k*_off_, leaving that target available for re-occupancy. Drug elimination (*k*_3_) acts on the free-ligand pool rather than on the complex (Section 4); it does not shorten an occupied interval but depletes the ligand available for rebinding, and so governs how often the occupied state is re-entered. Target degradation (*k*_deg_), where present, destroys the target rather than freeing it, ending the current interval and precluding any subsequent re-occupancy of that target. No single one of these constants determines cumulative occupancy; only EOT, which integrates their joint effect over time, does.

##### Remark 3

(When is Rebinding Negligible?). *The Copeland limit (k*_1_ = 0*) applies under specific conditions: when experiments are conducted under “infinite dilution” conditions where dissociated ligand is immediately removed or diluted below effective concentration; when the dissociation rate greatly exceeds the effective association rate (k*_2_ ≫ *k*_1_*b*(*t*)*) for all relevant times; or when drug elimination is much faster than rebinding (k*_3_ ≫ *k*_1_*b*(*t*)*), where k*_3_ *is the elimination rate discussed in Section 4. In typical in vivo settings, none of these conditions hold, and rebinding contributes significantly to target occupancy. This is why EOT provides a more complete measure of therapeutic duration than DTRT*.

We note that the relationship between the verbal motivation of DTRT and its mathematical definition 1*/k*_off_ has itself been a subject of discussion in the literature: Folmer [12] observed that the predictive value of DTRT often reduces to potency in contexts dominated by drug clearance, and Copeland’s own 2021 retrospective [15] acknowledges that the verbal framing of residence time has evolved to incorporate association kinetics and rebinding—factors that go beyond the original 1*/k*_off_ definition. EOT, as defined in Equation (8), provides the rigorous integral formulation that gives mathematical precision to these verbal intuitions, and reduces to DTRT = 1*/k*_off_ exactly in the limiting case where rebinding is absent (Section 2.4).

## 3 Ehrlich Occupancy Time in Closed Systems

We first analyze closed systems (no influx or efflux of species), which model typical in vitro binding assays. These systems approach equilibrium as *t* → ∞, allowing us to compute rEOT analytically via Proposition 1.

### 3.1 Fundamental Receptor-Ligand Binding

Consider the reversible bimolecular reaction:

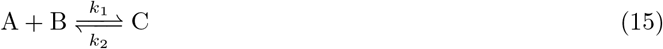

where *A* is the receptor (target), *B* is the drug (ligand), and *C* is the drug-receptor complex. Let *a*(*t*), *b*(*t*), and *c*(*t*) denote the concentrations of these species at time *t*, with initial conditions *a*(0) = *a*_0_, *b*(0) = *b*_0_ (units: M), and *c*(0) = 0 (no complex initially present).

Mass-action kinetics yields the system of ordinary differential equations:

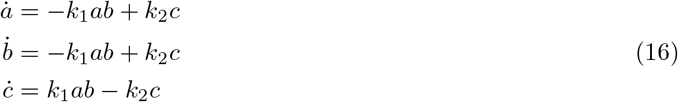

where *k*_1_ (units: M^*−*1^s^*−*1^) is the association rate constant and *k*_2_ (units: s^*−*1^) is the dissociation rate constant. The system satisfies two conservation laws:

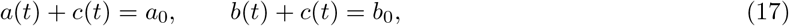

reflecting conservation of total receptor and total ligand, respectively. The fraction of occupied receptors at time *t* is *f* (*t*) = *c*(*t*)*/a*_0_.

#### 3.1.1 Equilibrium Analysis

At equilibrium, the forward and reverse reaction rates balance: *k*_1_*a*_*∞*_*b*_*∞*_ = *k*_2_*c*_*∞*_. Using the conservation laws (17) to substitute *a*_*∞*_ = *a*_0_ − *c*_*∞*_ and *b*_*∞*_ = *b*_0_ − *c*_*∞*_, we obtain:

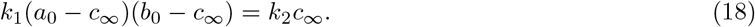

Expanding and rearranging yields a quadratic equation in *c*_*∞*_:

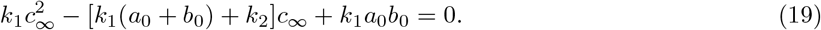

Applying the quadratic formula and selecting the physically meaningful root (satisfying 0 ≤ *c*_*∞*_ ≤ min(*a*_0_, *b*_0_)), we obtain after simplification:

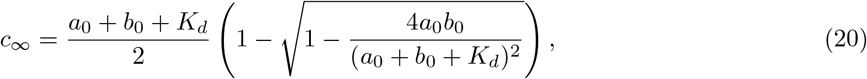

where *K*_*d*_ = *k*_2_*/k*_1_ is the **dissociation constant** (units: M). The equilibrium fraction of bound receptors is therefore:

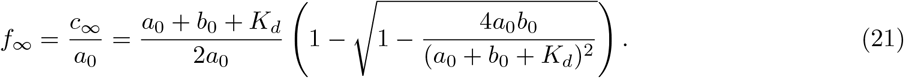

By Proposition 1, this equals the limiting relative EOT: rEOT(*T*) → *f*_*∞*_ as *T* → ∞.

#### 3.1.2 Pseudo-First-Order Approximation

The exact expression (21) is difficult to interpret. Considerable simplification occurs in the biologically common regime where the receptor concentration is much smaller than the ligand concentration: *a*_0_ ≪ *b*_0_. This **pseudo-first-order** (or **ligand excess**) approximation is standard in biochemical kinetics and applies when drug depletion due to target binding is negligible.

Under this assumption, define the dimensionless quantity:

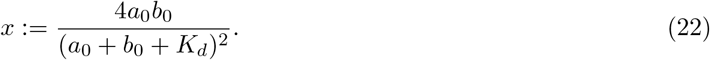

When *a*_0_ ≪ *b*_0_, we have *a*_0_ + *b*_0_ + *K*_*d*_ ≈ *b*_0_ + *K*_*d*_, so:

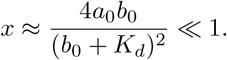

For small *x*, the Taylor expansion 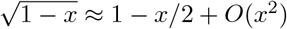 applies. Substituting into (21):

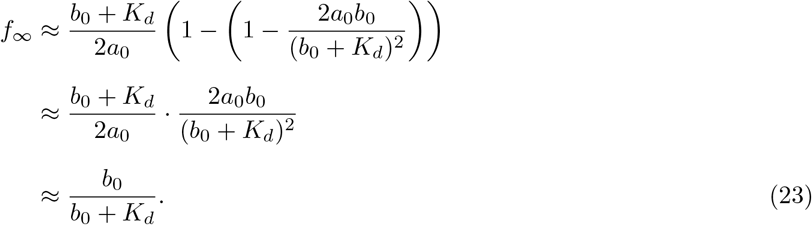

This yields the fundamental result for receptor-ligand binding under pseudo-first-order conditions:

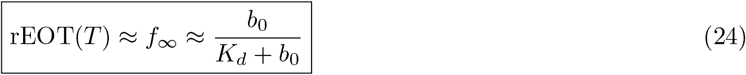

Note that *f*_*∞*_ *>* 0 whenever *b*_0_ *>* 0, which is guaranteed under the pseudo-first-order assumption (*b*_0_ ≫ *a*_0_ *>* 0). The degenerate case *f*_*∞*_ = 0 corresponds to the absence of drug (*b*_0_ = 0), which lies outside the scope of this approximation.

Equation (24) is the classical Langmuir isotherm, expressed here as the relative Ehrlich occupancy time. The isotherm exhibits three characteristic regimes: at low drug concentration (*b*_0_ ≪ *K*_*d*_), occupancy scales linearly with concentration as *f*_*∞*_ ≈ *b*_0_*/K*_*d*_; at the dissociation constant (*b*_0_ = *K*_*d*_), occupancy reaches its half-maximal value *f*_*∞*_ = 1*/*2; and at high drug concentration (*b*_0_ ≫ *K*_*d*_), occupancy approaches saturation with *f*_*∞*_ → 1.

##### Remark 4

(Validity of the Approximation). *The pseudo-first-order approximation requires a*_0_ ≪ *b*_0_, *ensuring that ligand depletion due to binding is negligible (i*.*e*., *b*(*t*) ≈ *b*_0_ *throughout the time course). This condition is typically satisfied in biochemical assays such as SPR and ITC that are designed with excess ligand, in therapeutic settings where drug concentrations (nM–µM) exceed target concentrations [1, 3], and in cellular assays where receptor density is low relative to extracellular drug. For systems where a*_0_ *and b*_0_ *are comparable, the full expression* (21) *must be used*.

#### 3.1.3 Biological Interpretation

Equation (24) reveals that relative EOT depends on both *k*_1_ and *k*_2_ through the dissociation constant *K*_*d*_ = *k*_2_*/k*_1_, not on *k*_2_ alone as in Copeland’s DTRT. This represents the correct mathematization of Ehrlich’s principle that drugs act when bound—cumulative occupancy at equilibrium is determined by *K*_*d*_, which emerges naturally from our framework.

While Copeland and coauthors [1] cite Ehrlich, their focus on *k*_2_ alone (DTRT = 1*/k*_2_) captures only single-event kinetics, missing the cumulative nature of target engagement that Ehrlich emphasized. The fundamental distinction, visualized in Figure 1, is that Copeland DTRT measures only the duration of individual binding events (panel D), while EOT captures the cumulative occupancy including rebinding (panel C). In closed systems where drug molecules remain available for rebinding after dissociation, EOT provides the complete measure of target engagement.

As illustrated in Figure 1C, every binding event—whether initial or subsequent rebinding—contributes to EOT. The two metrics coincide only when rebinding is absent, as shown in Section 2.4.

### 3.2 Induced Fit: Conformational Dynamics

Many drug-target interactions involve conformational changes upon binding [16, 17]. The drug may initially bind in one conformation, then induce (or select) a second conformation that alters the stability of the complex. This **induced fit** mechanism is represented by augmenting the basic binding reaction with an isomerization step:

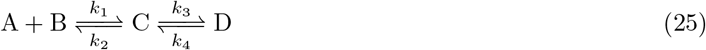

Here, *A* is the free receptor, *B* is the drug, *C* is the initial encounter complex, and *D* is the isomerized (conformationally changed) complex. The rate constants *k*_3_ and *k*_4_ govern the forward and reverse isomerization, respectively.

The corresponding mass-action kinetics are:

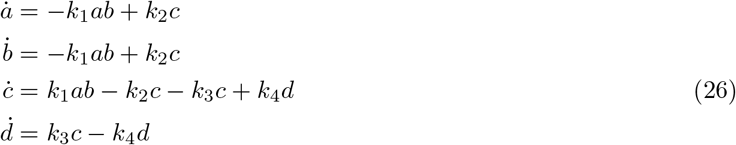

The conservation laws are:

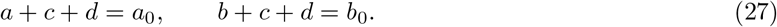

Since both *C* and *D* represent receptor-bound states, the fraction of occupied receptors is:

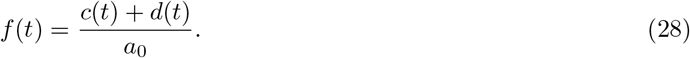

#### 3.2.1 Equilibrium Analysis

At equilibrium, both reactions are balanced:

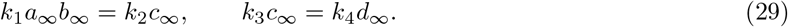

From the second condition, the isomerized complex concentration is proportional to the encounter complex:

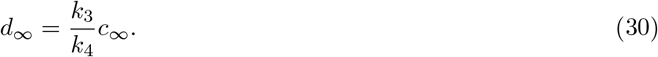

The total bound receptor concentration is therefore:

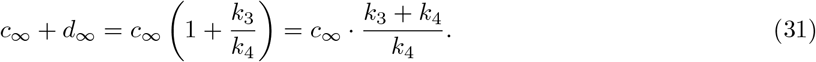

To simplify subsequent expressions, define:

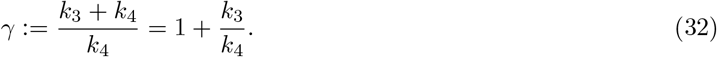

It is convenient to introduce the total bound concentration *s*_*∞*_:= *c*_*∞*_ + *d*_*∞*_ = *γc*_*∞*_. Then *c*_*∞*_ = *s*_*∞*_*/γ*, and the conservation laws (27) give

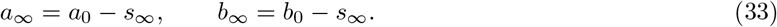

Substituting into the first equilibrium condition *k*_1_*a*_*∞*_*b*_*∞*_ = *k*_2_*c*_*∞*_ gives

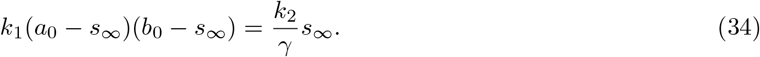

Using *K*_*d*_ = *k*_2_*/k*_1_, the total bound concentration therefore satisfies

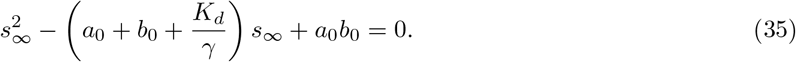

The physically meaningful solution (satisfying 0 ≤ *s*_*∞*_ ≤ min(*a*_0_, *b*_0_)) is

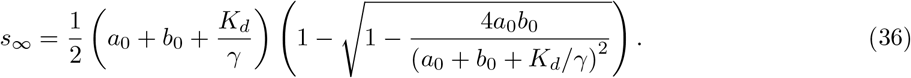

#### 3.2.2 Pseudo-First-Order Approximation

For the biologically common case *a*_0_ ≪ *b*_0_, we apply the same approximation strategy as in Section 3.1. Define

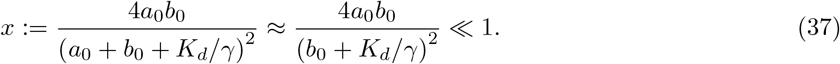

Using 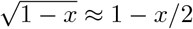 for small *x*,

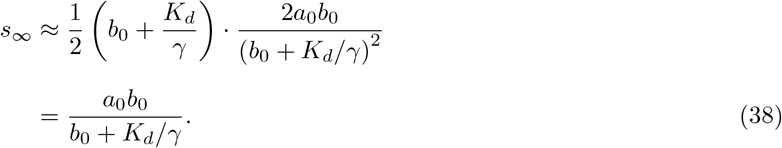

The equilibrium fraction of bound receptors is therefore

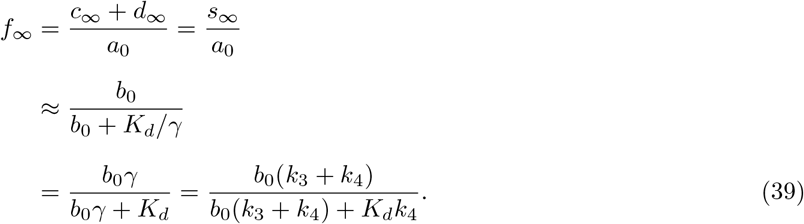

This can be rewritten more elegantly by defining an **effective dissociation constant**:

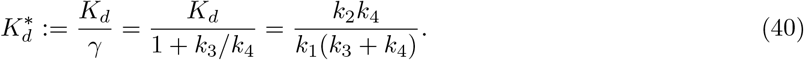

With this definition, the ligand-excess equilibrium occupancy takes the familiar Langmuir form:

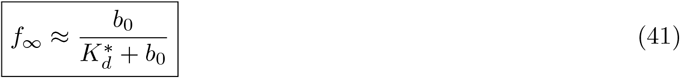

Since *γ* ≥1 (with equality only when *k*_3_ = 0), we have 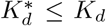. The induced fit mechanism effectively *increases* the apparent binding affinity.

#### 3.2.3 Clinical Examples of Induced Fit

The induced fit mechanism is not merely theoretical—it has been exploited successfully in drug design.

##### Example 2

(Type II Kinase Inhibitors). *Imatinib (Gleevec), used to treat chronic myeloid leukemia, binds to the BCR-ABL kinase in an inactive “DFG-out” conformation. Upon binding, the drug stabilizes a conformational state (complex D) that differs from the initial encounter complex (complex C). This conformational selection mechanism contributes to imatinib’s prolonged residence time* (~*30 minutes) despite moderate equilibrium affinity [8]. In our framework, a large k*_3_*/k*_4_ *ratio (favoring the DFG-out state) reduces* 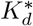 *below the intrinsic K*_*d*_, *enhancing target occupancy*.

##### Example 3

(GPCR Ligands). *Many G protein-coupled receptor (GPCR) drugs induce conformational changes that stabilize active or inactive receptor states. Tiotropium, a long-acting muscarinic antagonist for chronic obstructive pulmonary disease (COPD), exhibits slow dissociation (k*_2_ ~ 0.0002 *s*^*−*1^, *residence time* ~ *1*.*4 hours) partly due to conformational stabilization of an inactive receptor state. This allows once-daily dosing despite systemic elimination within hours*.

These examples demonstrate that optimizing the conformational landscape (large *k*_3_*/k*_4_) can achieve prolonged target occupancy even when initial binding (*k*_1_, *k*_2_) is not exceptionally favorable.

#### 3.2.4 Biological Interpretation

The conformational change acts as a kinetic “trap.” Fast forward isomerization (large *k*_3_) or slow reverse isomerization (small *k*_4_) increases the ratio *k*_3_*/k*_4_, thereby decreasing 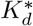 and increasing rEOT. This explains why induced fit inhibitors can achieve prolonged target occupancy even with moderate initial binding affinity.

To illustrate the magnitude of this effect, consider two limiting scenarios. In the absence of induced fit (*k*_3_ = 0), the effective dissociation constant reduces to 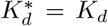, recovering simple binding behavior. With a strong induced fit (*k*_3_*/k*_4_ = 9), however, the effective dissociation constant becomes 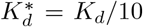, representing a 10-fold increase in apparent affinity purely from conformational trapping.

The framework quantifies how conformational dynamics contribute to residence time beyond simple binding kinetics.

##### Example 4

(Quantitative Impact). *Suppose K*_*d*_ = 1 *µM and b*_0_ = 1 *µM. Without induced fit*, 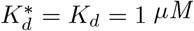, *yielding f*_*∞*_ = 1*/*(1 + 1) = 0.50. *With a strong induced fit (k*_3_*/k*_4_ = 9*), the effective dissociation constant drops to* 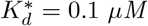, *yielding f*_*∞*_ = 1*/*(0.1 + 1) ≈ 0.91. *Thus, the induced fit mechanism increases target occupancy from 50% to 91%—an 82% relative increase—purely from conformational trapping, without changing the intrinsic binding rate constants k*_1_ *and k*_2_.

Our finding that induced fit mechanisms can dramatically extend occupancy time is consistent with Copeland’s recent acknowledgment of conformational dynamics in residence time [15]. However, our framework quantifies this effect through the modified dissociation constant 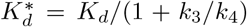, providing a direct mathematical relationship between conformational kinetics and occupancy time.

##### Remark 5

(Notation for *k*_3_). *In this section, k*_3_ *denotes the forward isomerization rate constant. In Section 4, k*_3_ *denotes the first-order drug-elimination rate constant. These parameters belong to distinct reaction mechanisms and are never used simultaneously. We retain the conventional sequential indexing of rate constants within each mechanism, while using the surrounding mechanism and descriptive terminology (forward isomerization rate versus elimination rate) to disambiguate them*.

## 4 Ehrlich Occupancy Time with Drug Removal

### 4.1 Biological Motivation

In vivo, drugs undergo continuous removal through metabolism, renal excretion, and other elimination processes. These pharmacokinetic factors fundamentally alter drug-target interactions by depleting the free drug pool available for binding. Unlike closed systems where EOT grows linearly with time, drug removal causes EOT to saturate at a finite limit EOT_*∞*_ as *T*→∞. Understanding this limit is crucial for predicting therapeutic duration and optimizing dosing regimens.

To illustrate the magnitude of these effects, consider a hypothetical drug with strong target binding (*K*_*d*_ = 1 nM, *k*_2_ = 0.001 s^*−*1^) administered intravenously at a dose achieving the initial plasma concentration *b*_0_ = 100 nM. With slow elimination (*k*_3_ = 0.0001 s^*−*1^, corresponding to *t*_1*/*2_ ~ 2 hours), the upper bound from Equation (58) gives 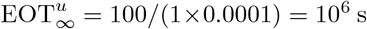, and target occupancy remains high over a 24-hour period. In stark contrast, with rapid elimination (*k*_3_ = 0.1 s^*−*1^, *t*_1*/*2_ ~ 7 seconds), the upper bound drops to 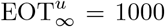 s—three orders of magnitude lower despite identical binding kinetics. Despite identical binding kinetics, rapid clearance limits effective target engagement to minutes rather than hours.

This 1000-fold difference cannot be predicted from binding affinity or Copeland DTRT alone. While it is qualitatively appreciated in drug discovery that rapid clearance can negate strong target binding [1, 3], our framework provides an explicit quantitative relationship between *k*_3_, *K*_*d*_, and cumulative occupancy time. This helps explain why some high-affinity compounds discovered in biochemical assays fail in vivo: excellent target binding is negated by poor pharmacokinetics. Our framework quantifies this interplay explicitly.

### 4.2 Model Formulation

Consider receptor-ligand binding coupled with first-order drug removal:

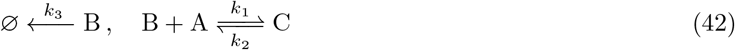

Here *k*_3_ ≡ *k*_elim_ represents the first-order drug-elimination rate constant, encompassing all removal processes (metabolism, excretion, distribution out of the compartment). The mass-action equations are:

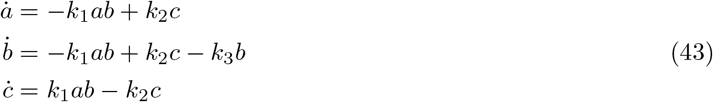

with initial conditions *a*(0) = *a*_0_, *b*(0) = *b*_0_, *c*(0) = 0. The conservation law *a* + *c* = *a*_0_ still holds (receptor is conserved), but total ligand is no longer conserved due to elimination.

Unlike closed systems, *b*(*t*) → 0 as *t* → ∞ due to removal, driving the system toward complete dissociation (*c*(*t*) → 0). Consequently, EOT(*T*) approaches a finite limit EOT_*∞*_ rather than growing indefinitely.

#### 4.2.1 Comparison with the DTRT Approach

To illustrate the limitations of Copeland’s DTRT approach in this context, consider what happens if we apply the DTRT methodology to our system. Following Copeland et al. [1], one would wait for the system to reach a quasi-steady state (*a*^*\**^, *b*^*\**^, *c*^*\**^) with positive entries, then “switch off” ligand association (mimicking infinite dilution). The remaining dynamics are:

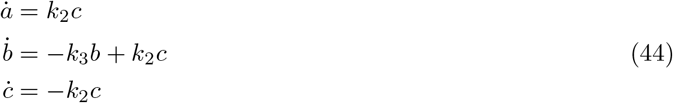

This linear system has eigenvalues 0, −*k*_3_, and −*k*_2_. While the complete system dynamics are governed by the smaller of *k*_2_ and *k*_3_, the DTRT approach focuses solely on the third equation, concluding that *k*_2_ alone determines residence time (DTRT = 1*/k*_2_).

This analysis reveals three critical limitations of the DTRT approach. First, DTRT neglects the elimination rate *k*_3_ entirely, yet when *k*_3_ *> k*_2_, drug elimination dominates the long-term behavior and determines the duration of target engagement. Second, because it is computed under the infinite-dilution assumption, DTRT reflects only a single binding event and not the additional occupancy that rebinding—the reassociation of dissociated drug molecules with nearby targets—contributes in vivo. Third, the single-parameter metric (1*/k*_2_) fundamentally cannot capture the multi-parameter dependencies revealed by our EOT framework, where both binding kinetics and pharmacokinetics jointly determine cumulative occupancy.

Our EOT analysis shows that both *k*_2_ and *k*_3_ contribute to determining occupancy time, with their relative importance determined by the ratio *k*_3_*/k*_2_.

### 4.3 Pseudo-First-Order Approximation

The system (43) does not admit a closed-form solution in general. However, under the assumption of excess ligand (*a*_0_ ≪ *b*_0_), we can derive a tractable approximation. This assumption is standard in many experimental and therapeutic settings: therapeutic plasma concentrations typically range from nanomolar to micromolar, while many drug targets exist at substantially lower concentrations (e.g., intracellular kinases are often present at only 1–100 nM)—provided that the drug penetrates the relevant cellular compartment such that free intracellular drug concentrations are comparable to plasma concentrations; cases where active efflux or poor membrane permeability substantially reduce intracellular exposure require cellular measurements of *b*_0_ rather than plasma values [11]—and biochemical binding assays such as SPR and ITC are typically designed with excess ligand to ensure measurable signals. The approximation is valid whenever drug depletion due to target binding is negligible, i.e., when the amount of drug bound to targets remains small compared to the total amount of drug present.

#### 4.3.1 Derivation of the Reduced System

The small parameter governing the pseudo-first-order regime is the dimensionless concentration ratio

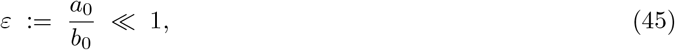

which measures the degree to which drug depletion due to target binding is negligible. To apply perturbation theory systematically, we introduce *O*(1) dimensionless scaled variables

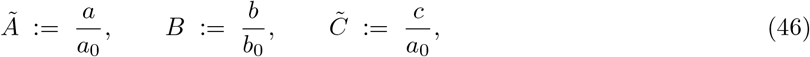

with initial conditions *Ã* (0) = *B*(0) = 1 and 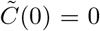. By construction, all three variables are *O*(1) for all *t* ≥ 0. The conservation law *a* + *c* = *a*_0_ becomes 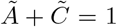. Substituting 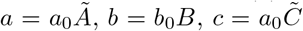 into system (43) and using *a*_0_ = *εb*_0_ yields the *ε*-parameterized dimensionless system:

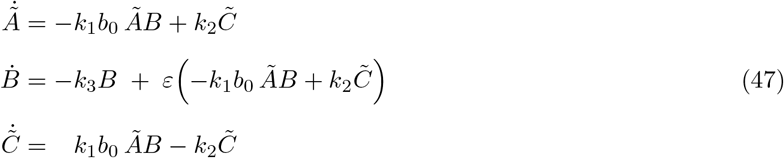

The critical observation is that all three scaled variables are *O*(1), and the coupling of the drug equation 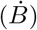 to the receptor and complex dynamics appears *only* at *O*(*ε*). Setting *ε* = 0 in (47) therefore decouples the drug equation at leading order:

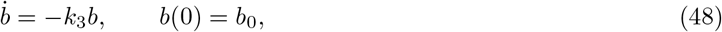

(equivalently, the *O*(1) equation 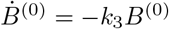, *B*^(0)^(0) = 1), with solution:

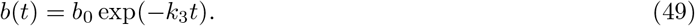

Substituting the leading-order approximation *B*^(0)^(*t*) = exp(−*k*_3_*t*) into the 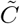 equation of (47), invoking the conservation relation 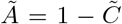, and returning to dimensional variables via 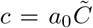, yields the *O*(1) approximation for the complex concentration:

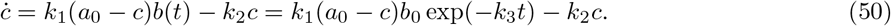

Rearranging:

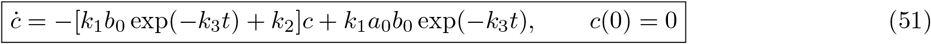

This is a linear, non-autonomous first-order ODE. The Ehrlich occupancy time is:

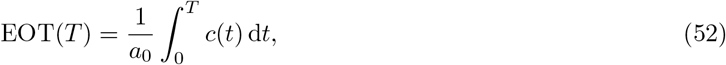

and we seek EOT_*∞*_ = lim_*T→∞*_ EOT(*T*).

#### 4.3.2 Solution Structure

The solution to (51) can be expressed using variation of constants. The homogeneous equation

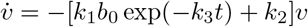

has solution:

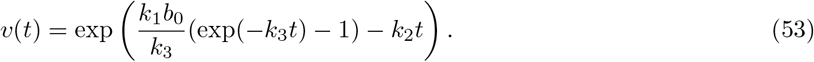

While this leads to an exact integral representation of *c*(*t*), the resulting integrals generally cannot be evaluated in closed form. Instead, we derive rigorous upper and lower bounds using comparison principles for differential equations [18].

### 4.4 Analytical Bounds for EOT_*∞*_

#### 4.4.1 Upper Bound

From Equation (51), since *c* ≥ 0 and exp(−*k*_3_*t*) ≤ 1 for *t* ≥ 0, we have the differential inequality:

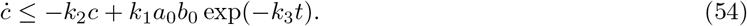

With the integrating factor *µ*(*t*) = exp(*k*_2_*t*) we get

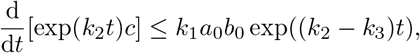

and integrating from 0 to *t* (assuming *k*_2_ ≠ *k*_3_; the borderline case yields the same final bounds) we find

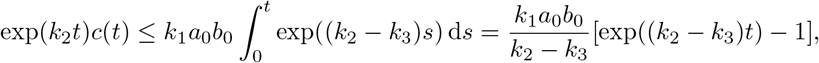

therefore:

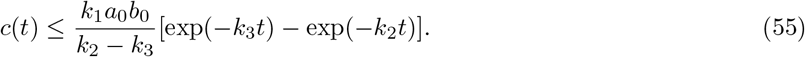

To obtain the bound on EOT_*∞*_, we integrate:

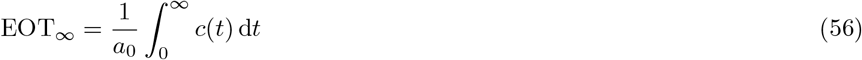

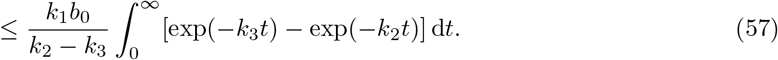

Using the standard result 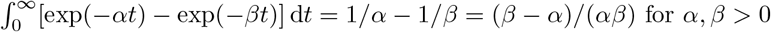:

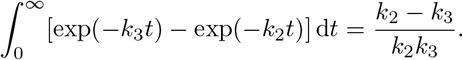

Substituting:

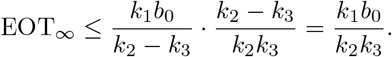

Expressing in terms of *K*_*d*_ = *k*_2_*/k*_1_:

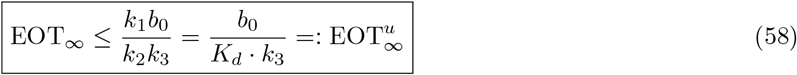

##### Remark 6

*Our original argument (here and in the following subsection) was more roundabout, using differential inequalities with the comparison equation:*

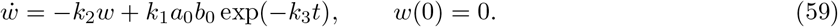

*As noted by a reviewer, invoking the comparison principle is not strictly necessary here, and the argument can be streamlined. Comparison principles generally extend to larger classes of equations, and will be used in Appendix A below, in the derivation of different estimates*.

#### 4.4.2 Lower Bound

From Equation (51), since exp(−*k*_3_*t*) ≤ 1 and *c* ≥ 0:

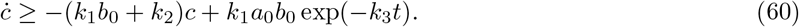

Using the integrating factor *µ*(*t*) = exp((*k*_1_*b*_0_ + *k*_2_)*t*):

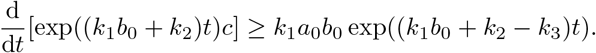

Integrating from 0 to *t*:

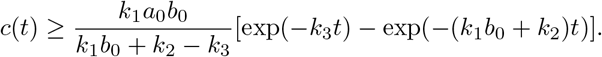

Integrating over [0, ∞):

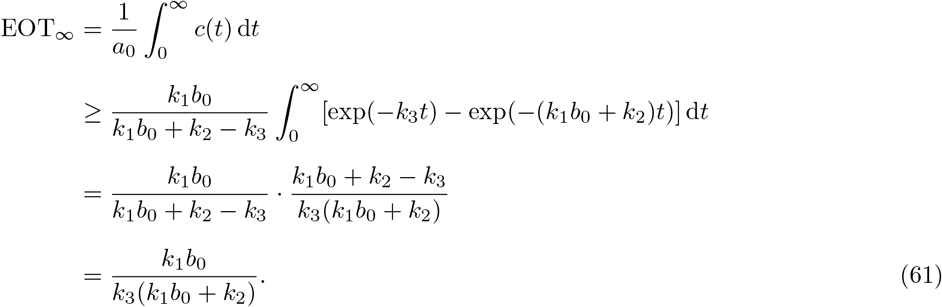

Expressing in terms of *K*_*d*_:

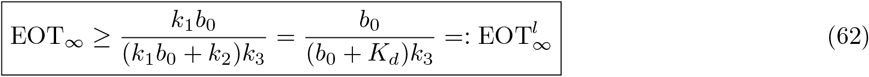

#### 4.4.3 Summary of Bounds

Combining equations (58) and (62):

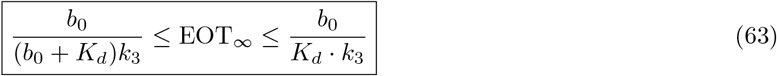

The relationship between the bounds can be expressed as:

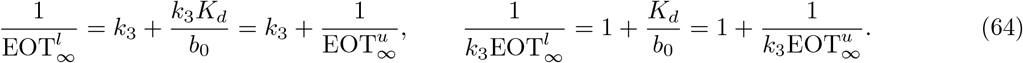

The ratio of upper to lower bounds is:

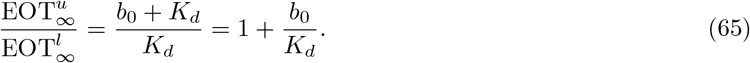

#### 4.4.4 Sharpness of the Bounds

The bounds (63) are tightest when *b*_0_ ≪ *K*_*d*_ (low drug concentration relative to affinity), where 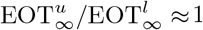. The bounds diverge when *b*_0_ ≫ *K*_*d*_ (high concentration), but in this regime both bounds predict large EOT_*∞*_, so the qualitative conclusion (prolonged occupancy) remains valid.

More precisely, we can compare two sets of bounds. The *standard bounds* (equations (58)–(62)), derived above using comparison with constant-coefficient equations, are complemented by *exponential bounds* derived using variation of constants (Appendix A). The exponential bounds take the form:

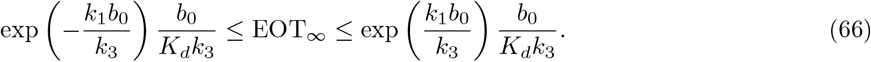

The standard upper bound is always tighter than the exponential upper bound. For the lower bounds, the standard bound is tighter when:

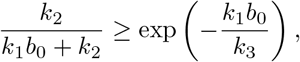

which is satisfied whenever *k*_2_ *> k*_3_. Since the biologically common regime has dissociation faster than elimination (*k*_2_ *> k*_3_), the standard bounds (63) are typically the most useful.

### 4.5 Biological Interpretation

#### 4.5.1 Effect of Removal Rate *k*_3_

Both bounds are inversely proportional to *k*_3_. Faster drug elimination (larger *k*_3_) proportionally reduces EOT_*∞*_. While it is broadly appreciated in drug discovery that pharmacokinetics and pharmacodynamics must both be optimized [2, 3], our bounds make this trade-off mathematically precise: Equation (63) shows explicitly that *k*_3_ and *K*_*d*_ enter multiplicatively, so that no amount of binding affinity optimization can compensate for unlimited increases in elimination rate. This reveals a fundamental trade-off: even drugs with very slow dissociation (small *k*_2_, large Copeland DTRT) achieve only limited target occupancy if eliminated rapidly. For example, if *k*_3_ ≫ *k*_2_, removal dominates and EOT_*∞*_ is primarily limited by pharmacokinetics rather than binding kinetics. From Equation (58), doubling *k*_3_ halves the maximum achievable EOT_*∞*_ regardless of binding affinity.

##### Rebinding in the presence of elimination

The interplay between rebinding and drug removal deserves special attention. In a closed system (Section 3), a dissociated drug molecule remains available indefinitely for rebinding, leading to multiple occupancy cycles as depicted in Figure 1C. With drug removal, however, each dissociation event is potentially final: the probability that a dissociated drug molecule rebinds before being eliminated is approximately *k*_1_*b*(*t*)*/*(*k*_1_*b*(*t*) + *k*_3_).

As drug concentration *b*(*t*) = *b*_0_ exp(−*k*_3_*t*) declines exponentially, this probability decreases over time. Early in the time course, when *b*(*t*) ≈ *b*_0_, rebinding remains likely and the system exhibits dynamics similar to the closed case. At later times, when *b*(*t*) ≪ *K*_*d*_, rebinding becomes rare, and the system approaches the Copeland limit where each receptor experiences at most one binding event.

This temporal evolution from a rebinding-dominated regime to a single-binding regime explains why our bounds depend on both *K*_*d*_ and *k*_3_: they capture the full spectrum of behaviors across the entire time course.

#### 4.5.2 Effect of Dissociation Constant *K*_*d*_

The upper bound 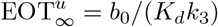 shows that tighter binding (smaller *K*_*d*_) increases occupancy time, as expected. However, the lower bound 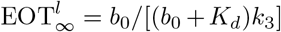 reveals more nuanced behavior depending on drug concentration relative to affinity. Under saturating conditions (*b*_0_ ≫ *K*_*d*_), the lower bound simplifies to 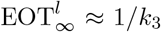, becoming independent of binding affinity—once targets are saturated, occupancy duration is determined entirely by how long drug remains available. Conversely, under subsaturating conditions (*b*_0_ ≪ *K*_*d*_), the lower bound becomes 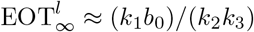, showing that both association and dissociation rates contribute to determining occupancy time.

#### 4.5.3 Parameter Regimes

We identify three key regimes based on the ratio *k*_3_*/k*_2_:

1. *k*_3_ ≪ *k*_2_ **(slow removal):** Drug clearance is slower than unbinding. EOT_*∞*_ is primarily determined by binding kinetics, and the system behaves similarly to a closed system over therapeutically relevant timescales. The upper bound dominates: EOT_*∞*_ ≈ *b*_0_*/*(*K*_*d*_*k*_3_).
2. *k*_3_ ≈ *k*_2_ **(comparable rates):** Both binding kinetics and pharmacokinetics contribute significantly. The bounds (63) provide tight estimates. Optimization requires balancing affinity and elimination rate.
3. *k*_3_ ≫ *k*_2_ **(fast removal):** Drug is eliminated before significant binding equilibration. The lower bound dominates: EOT_*∞*_ ≈ *b*_0_*/*[(*b*_0_ + *K*_*d*_)*k*_3_]. In this regime, improving binding affinity has limited benefit unless *k*_3_ is also reduced (e.g., via formulation changes, prodrugs, or reducing first-pass metabolism).

##### Decision framework for drug optimization

The bounds (63) provide actionable guidance for medicinal chemistry optimization, as summarized in Table 2. Appendix B provides detailed experimental protocols for measuring the required parameters and a case study illustrating this optimization workflow.

**Table 1.** Glossary of key terms and parameters.

| Term/Symbol | Definition |
| --- | --- |
| EOT | Ehrlich Occupancy Time: cumulative duration target is occupied, including all binding/unbinding cycles |
| rEOT | Relative EOT: $EOT(T)/T$, converges to equilibrium occupancy fraction |
| DTRT | Drug-Target Residence Time (Copeland): $1/k_{\text{off}}$, duration of single binding event |
| $k_1$ (or $k_{\text{on}}$) | Association rate constant ($M^{-1}s^{-1}$): how fast drug binds to free target |
| $k_2$ (or $k_{\text{off}}$) | Dissociation rate constant ($s^{-1}$): how fast drug-target complex dissociates |
| $k_3$ (Section 3.2) | Forward isomerization rate constant ($s^{-1}$) in the induced-fit mechanism |
| $k_3 \equiv k_{\text{elim}}$ (Section 4) | First-order drug-elimination rate constant ($s^{-1}$) in the drug-removal mechanism |
| $K_d$ | Dissociation constant ($k_2/k_1$, units M): lower $K_d$ = tighter binding |
| $f(t)$ | Fractional target occupancy at time $t$: fraction of targets bound by drug |
| $a_0, b_0$ | Initial concentrations of target and drug, respectively |
| Rebinding | Drug molecule dissociates then reassociates before elimination |
| Induced fit | Conformational change in target upon drug binding (mechanism in equation (25)) |
| Pseudo-first-order | Approximation valid when target concentration ≪ drug concentration |

**Table 2.** Decision framework for drug optimization based on EOT analysis.

| Observation | Diagnosis | Optimization Strategy |
| --- | --- | --- |
| Low cellular potency despite good biochemical $K_d$ | Likely $k_3 \gg k_2$: fast removal limits occupancy | Reduce $k_3$ by minimizing metabolic soft spots or blocking efflux transporters; increase $k_2^{-1}$ by optimizing for slower dissociation |
| Strong biochemical but weak in vivo efficacy | $k_3$ limits $\text{EOT}_\infty$ despite good $K_d$ | Consider formulation strategies (sustained release), prodrug approaches, or combination with metabolic inhibitors to reduce effective $k_3$ |
| Good potency but short duration of action | Slow off-rate ($k_2^{-1}$) not compensating for large $k_3$ | Prioritize compounds with ultra-slow $k_2$ or explore covalent inhibitors (irreversible binding) |
| Adequate efficacy but requiring frequent dosing | $\text{EOT}_\infty$ sufficient but duration limited by $k_3$ | Optimize PK profile rather than binding kinetics; consider depot formulations |

### 4.6 Comparison with Copeland DTRT

Copeland’s framework defines DTRT = 1*/k*_2_, independent of *k*_3_. This predicts that two drugs with identical *k*_2_ but different *k*_3_ values would have identical residence times—clearly incorrect in vivo. Our EOT framework correctly predicts that faster elimination (larger *k*_3_) reduces occupancy time proportionally, regardless of *k*_2_.

The conceptual limitation of Copeland DTRT becomes particularly apparent in the drug removal scenario. As shown in Figure 1D, Copeland’s approach counts only the duration of the first binding event for each receptor. However, in the presence of drug removal, whether a receptor experiences subsequent binding events (Figure 1C) depends critically on *k*_3_: fast elimination reduces the likelihood of rebinding by depleting available drug molecules. Since Copeland DTRT ignores rebinding entirely by definition, it cannot capture this pharmacokinetic effect.

#### Example 5

(Comparison of Two Drugs). *Consider two drugs with identical association and dissociation kinetics (k*_1_ = 10^6^ *M*^*−*1^*s*^*−*1^, *k*_2_ = 0.1 *s*^*−*1^*) but different elimination rates: Drug A has slow elimination (k*_3_ = 0.01 *s*^*−*1^*) while Drug B has rapid elimination (k*_3_ = 1.0 *s*^*−*1^*). Both drugs have identical K*_*d*_ = 0.1 *µM and Copeland DTRT of 10 s. However, applying equation* (58), *Drug A achieves* 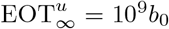 *while Drug B achieves only* 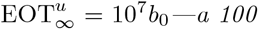 *-fold difference in cumulative target occupancy that is entirely invisible to the Copeland framework but crucial for predicting in vivo efficacy*.

Table 3 provides additional comparative examples that illustrate how Copeland DTRT can be misleading when elimination rates differ.

**Table 3.**
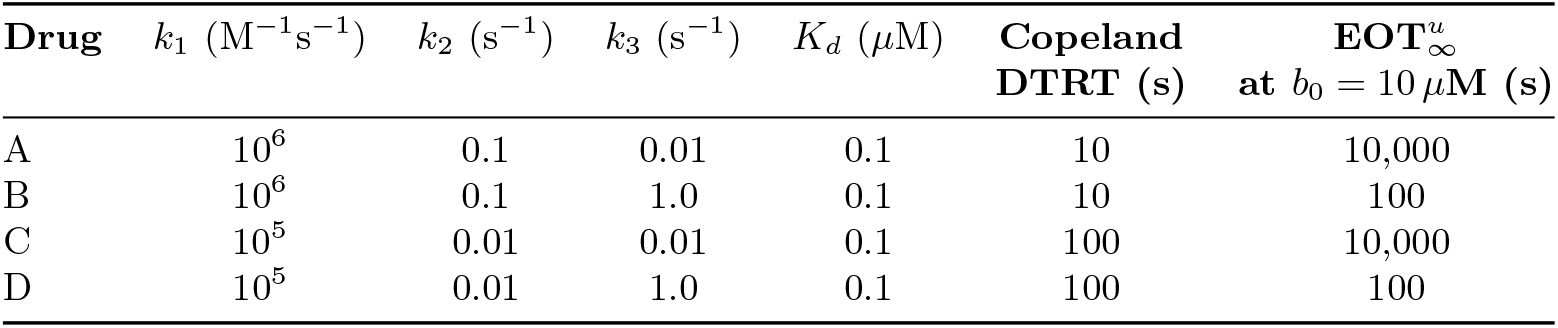
Example calculations comparing Copeland DTRT and EOT_*∞*_ for hypothetical drugs. *Interpretation:* Drugs A and B have identical Copeland DTRT but 100-fold different EOT_*∞*_ due to elimination rate differences. Drugs A and C have identical EOT_*∞*_ despite 10-fold different Copeland DTRT. This demonstrates that Copeland DTRT alone cannot predict in vivo occupancy time when drug elimination is significant.

### 4.7 Numerical Validation

Figures 2 and 3 present extensive numerical simulations validating our theoretical bounds across a comprehensive parameter space. We solve the full system (43) numerically for over 1,600 parameter combinations and compare with bounds (63).

**Figure 2.**
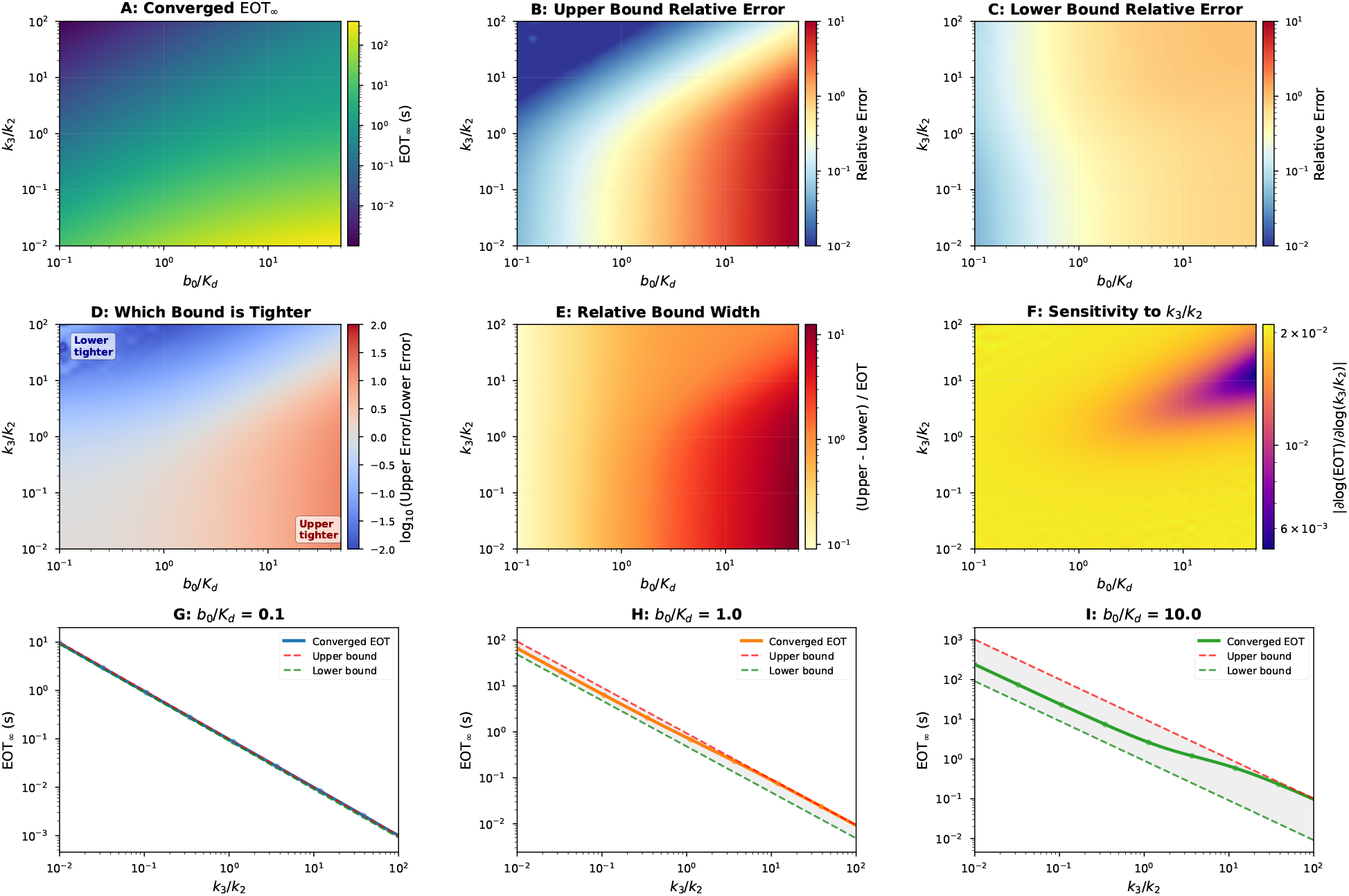
High-resolution analysis of EOT bounds across parameter space. **(A)** Converged EOT_*∞*_ values showing the full landscape of occupancy times across four orders of magnitude in *k*_3_*/k*_2_ and nearly two orders in *b*_0_*/K*_*d*_. **(B)** Upper bound relative error, demonstrating the bound is tightest (error *<* 0.1) when *b*_0_ ≪ *K*_*d*_ (low drug concentration regime). **(C)** Lower bound relative error, demonstrating the bound is tightest when *b*_0_ ≫ *K*_*d*_ (saturation regime). **(D)** Bound tightness comparison map, where blue regions indicate the lower bound is closer to the exact solution and red regions indicate the upper bound is closer. **(E)** Relative bound width 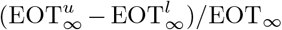, showing the bounds are most precise at extreme *k*_3_*/k*_2_ values. **(F)** Sensitivity analysis showing |*∂* log(EOT)*/∂* log(*k*_3_*/k*_2_)|, identifying regions of high parameter sensitivity. **(G-I)** Line plots of EOT_*∞*_ vs *k*_3_*/k*_2_ for *b*_0_*/K*_*d*_ = {0.1, 1.0, 10.0} respectively, with computed data points (dots, every 5th point shown), smooth interpolated curves (solid lines), upper bounds (red dashed), lower bounds (green dashed), and shaded gray regions indicating the valid bound range. The bounds provide reliable estimates across six orders of magnitude in both parameters. All simulations maintain pseudo-first-order validity (*a*_0_*/b*_0_ ≤ 0.01) to ensure bound accuracy. The heatmaps are computed on a 40×40 grid and displayed with 200×200 bicubic interpolation for smooth visualization.

**Figure 3.**
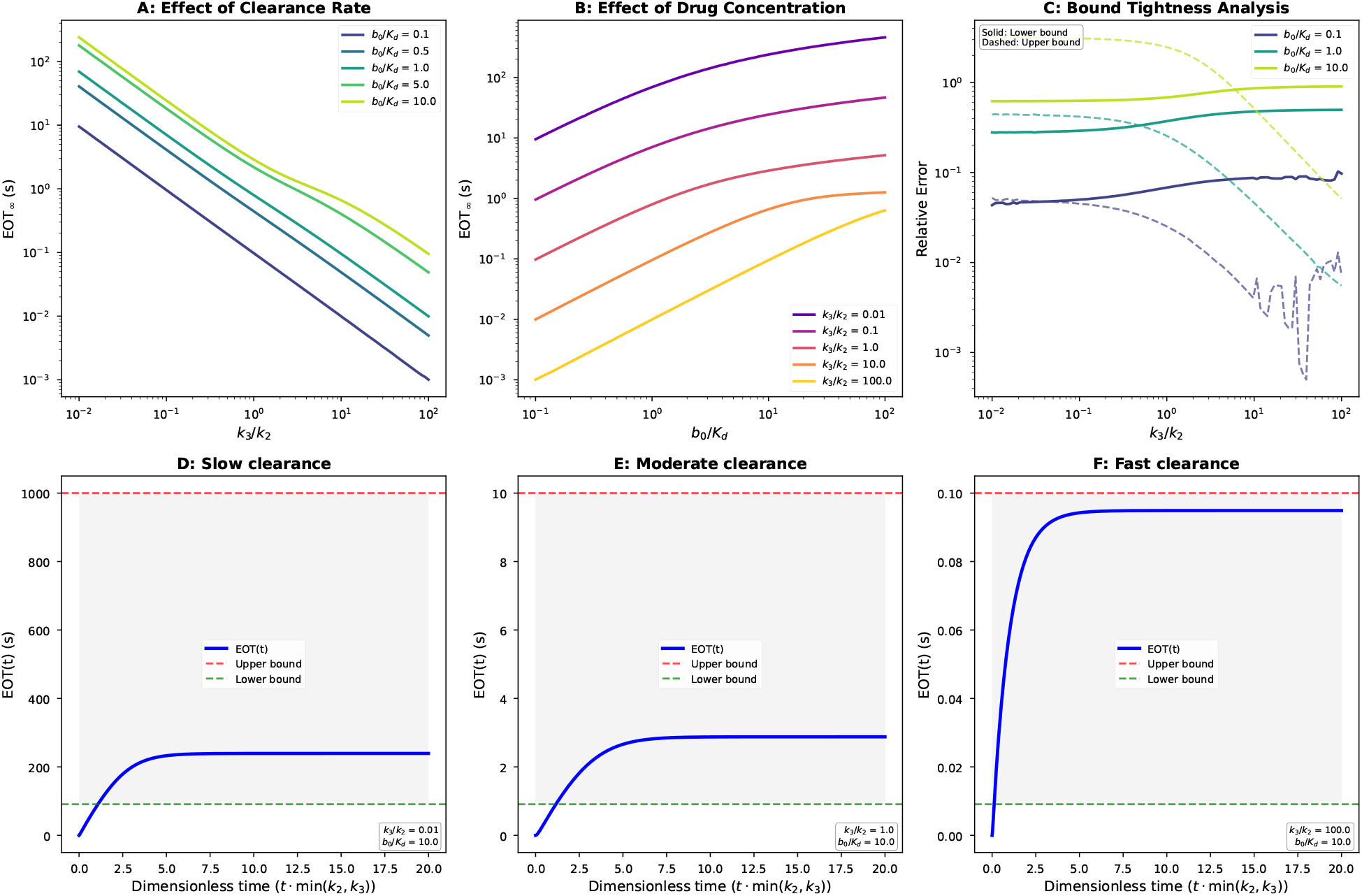
Systematic analysis of EOT behavior and bound quality. **(A)** Effect of clearance rate: EOT_*∞*_ vs *k*_3_*/k*_2_ for different drug concentrations (*b*_0_*/K*_*d*_ = 0.1 to 10), showing the transition from dissociation-limited (left) to clearance-limited (right) regimes. **(B)** Effect of drug concentration: EOT_*∞*_ vs *b*_0_*/K*_*d*_ for different *k*_3_*/k*_2_ ratios. **(C)** Bound tightness analysis showing relative error of upper (dashed) and lower (solid) bounds across parameter space. **(D–F)** Time evolution of EOT(*t*) approaching EOT_*∞*_ for three representative clearance rates at fixed *b*_0_*/K*_*d*_ = 10: (D) slow clearance (*k*_3_*/k*_2_ = 0.01), (E) moderate clearance (*k*_3_*/k*_2_ = 1), and (F) fast clearance (*k*_3_*/k*_2_ = 100). Blue curves show numerical EOT(*t*), horizontal dashed lines indicate analytical bounds (red: upper, green: lower), and gray shading shows the bound region. Panels D–F maintain pseudo-first-order validity (*a*_0_*/b*_0_ = 0.01), ensuring the analytical bounds are accurate.

#### 4.7.1 Computational Approach

The simulations span *k*_3_*/k*_2_ ratios from 0.01 to 100 (four orders of magnitude) and *b*_0_*/K*_*d*_ ratios from 0.1 to 50 (sub-saturating to saturating regimes). Computations were performed on a 40 × 40 parameter grid with bicubic interpolation to 200 × 200 for visualization. Throughout, validity of the pseudo-first-order approxi-mation was ensured by maintaining *a*_0_*/b*_0_ ≤ 0.01. For the converged EOT_*∞*_ calculations shown in Figures 2 and 3A–C, numerical solutions of the reduced pseudo-first-order Equation (51) were obtained in Python using `scipy.integrate.odeint` with relative and absolute tolerances set to 10^*−*10^ and 10^*−*12^, respectively [19]. For the illustrative time-course panels in Figure 3D–F, we used `scipy.integrate.solve ivp` with the stiff `Radau` solver and the same tolerances. The integral defining EOT_*∞*_ was then computed from the numerical solution *c*(*t*) via the trapezoidal rule (82), integrating until the running integral changed by less than 10^*−*8^ per unit time.

All scripts used to generate Figures 2–5 are publicly available at https://github.com/schnell-lab/ehrlich-occupancy-time, together with a `README` specifying the software environment and step-by-step instructions for reproducing each figure.

#### 4.7.2 Key Observations

The numerical results (Figure 2) confirm that our analytical bounds bracket the true EOT_*∞*_ across all parameter regimes tested. The upper bound is tightest when *b*_0_ ≪ *K*_*d*_ (panels B and D, blue regions), while the lower bound is tightest when *b*_0_ ≫ *K*_*d*_ (panels C and D, red regions). Both bounds converge in extreme regimes where either *k*_3_ ≪ *k*_2_ or *k*_3_ ≫*k*_2_ (panel E), and overall the bounds remain valid across six orders of magnitude in parameter space.

Figure 3 further demonstrates the time evolution and parameter dependencies, showing that EOT(*t*) approaches EOT_*∞*_ within the analytical bounds when the pseudo-first-order approximation is satisfied (panels D–F).

## 5 Discussion

We have presented a mathematically rigorous framework for *Ehrlich occupancy time* (EOT) that unifies classical residence-time concepts with dynamic ligand availability and conformational change. The central definition, 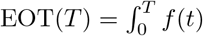 where *f* (*t*) is the fraction of occupied receptors at time *t*, captures cumulative occupancy including rebinding events and drug elimination effects—features absent from Copeland’s drug-target residence time (DTRT = 1*/k*_2_), which considers only the lifetime of a single binding event.

In closed systems approaching equilibrium, we proved that relative EOT converges to the equilibrium occupancy fraction *f*_*∞*_; under ligand-excess conditions this reduces to *f*_*∞*_ ≈ *b*_0_*/*(*K*_*d*_ + *b*_0_), depending on the dissociation constant *K*_d_ = *k*_2_*/k*_1_ rather than on *k*_2_ alone. For induced-fit mechanisms, conformational trapping reduces the effective dissociation constant to 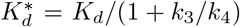, providing a quantitative link between conformational kinetics and prolonged occupancy.

In open systems with first-order drug elimination at rate *k*_3_, we derived rigorous bounds:

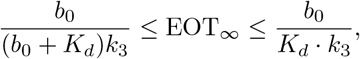

demonstrating that long-time occupancy is governed jointly by binding affinity and elimination rate. The numerical validation (Figures 2 and 3) confirms that these bounds bracket the true EOT_*∞*_ across six orders of magnitude in parameter space, with the bounds tightest when *b*_0_ ≪ *K*_*d*_ (upper bound) or *b*_0_ ≫*K*_*d*_ (lower bound).

### 5.1 Relation to Existing Residence-Time Metrics

Our framework contains Copeland’s DTRT as the limiting case in which rebinding is absent—for example, under infinite dilution or sufficiently rapid post-dissociation ligand removal. In closed systems with persistent ligand, EOT generally exceeds DTRT because rebinding contributes additional occupancy time. Outside this regime—particularly when exposure decays on the same timescale as dissociation, or when conformational gating is operative—DTRT misestimates the pharmacologically relevant occupancy time. In contrast, EOT is *exposure-aware* and *mechanism-aware*, incorporating association, dissociation, rebinding, conformational isomerization, and drug elimination within a single analytic scheme. Table 4 summarizes the key differences.

**Table 4.** Comparison of residence time definitions.

| Feature | Copeland DTRT | Ehrlich EOT (This Work) |
| --- | --- | --- |
| Mathematical definition | $1/k_{\text{off}}$ | $\int_0^T f(t) dt$ |
| Depends on $k_{\text{on}}$ | No | Yes |
| Includes rebinding | No<sup>a</sup> | Yes |
| Accounts for drug removal | No | Yes |
| Handles conformational change | Limited<sup>b</sup> | Yes (eq. (41)) |
| Closed-form solution | Always | Closed systems only |
| Bounds for open systems | N/A | Equations (63) |
| Valid when $k_3 > k_2$ | No | Yes |
| Use for in vivo occupancy prediction | Limited on its own | Broader, exposure-aware |

<sup>a</sup>Recent work acknowledges rebinding importance [15] but does not quantify it.
<sup>b</sup>An expression for $k_{\text{off}}$ is indicated in [1]; it does not incorporate rebinding.

Recent developments in residence-time theory have moved toward acknowledging these complexities. Copeland’s 2021 review [15] recognizes that intracellular residence time depends on the association rate constant and that cellular residence times are often prolonged by rebinding cycles. Our EOT framework quantifies these effects explicitly through the concentration-dependent terms in our bounds. Similarly, Folmer’s critique [12] that residence time provides a limited view of binding kinetics is addressed by our multi-parameter approach incorporating *k*_1_, *k*_2_, *k*_3_, and *b*_0_.

### 5.2 Implications for Drug Design

A central parameter emerging from our analysis is the dimensionless ratio *k*_3_*/k*_2_. When this ratio is small (drug persists longer than the complex lifetime), extending residence time by reducing *k*_2_ materially increases cumulative occupancy. When *k*_3_*/k*_2_ is moderate to large (exposure decays quickly relative to dissociation), modulating exposure—via formulation, dosing schedule, or distribution to protected compartments—dominates efficacy. The EOT_*∞*_-versus-(*k*_3_*/k*_2_) curves (Figure 2, panels G–I) quantify how much improvement is available from each optimization strategy.

These findings have several practical implications. First, binding affinity alone is insufficient for predicting efficacy: two drugs with identical *K*_*d*_ can have vastly different EOT values if their individual rate constants (*k*_1_, *k*_2_) or elimination rates (*k*_3_) differ. Second, pharmacokinetics and pharmacodynamics are inseparable when predicting target occupancy—the bounds show that *k*_3_ (a PK parameter) and *K*_*d*_ (a PD parameter) contribute multiplicatively, so optimizing one while neglecting the other may yield no net benefit. Third, for targets exhibiting induced fit, compounds triggering favorable conformational changes can achieve prolonged occupancy through kinetic trapping even with moderate initial binding affinity.

For experimentalists, the EOT framework provides concrete, testable targets: measuring *k*_1_ and *k*_2_ under conditions representing the biological compartment, characterizing the effective elimination rate *k*_3_, and identifying induced-fit signatures from multiphasic dissociation kinetics. Appendix B provides detailed protocols for these measurements along with a case study workflow for drug optimization.

### 5.3 Common Misconceptions

Based on discussions with experimental collaborators, we address three frequent misunderstandings.

First, *high affinity does not automatically imply long occupancy time*. While low *K*_*d*_ increases equilibrium occupancy, cumulative EOT depends additionally on elimination rate. A high-affinity drug can have short EOT if eliminated rapidly, because the system never reaches equilibrium before clearance. This explains why some high-affinity compounds fail in vivo despite excellent biochemical profiles.

Second, *Copeland residence time does not predict in vivo duration* in general. DTRT = 1*/k*_2_ measures the lifetime of a single binding event, whereas in vivo duration depends on EOT, which includes rebinding and elimination. As shown in Table 3, drugs with identical DTRT can have 100-fold different EOT due to differences in *k*_3_.

Third, *slow off-rate is not always beneficial*. Slow *k*_2_ helps only if drug remains available for binding; when *k*_3_ ≫ *k*_2_, elimination dominates and further reducing *k*_2_ provides diminishing returns. Moreover, extremely slow dissociation can cause on-target toxicity, reduce dose titratability, slow onset of action, or provide no benefit when target turnover is rapid. The optimal strategy is to match residence time to the desired therapeutic duration, considering target turnover and dosing frequency.

### 5.4 Limitations and Future Directions

We have idealized several processes as first-order (clearance, turnover) and treated induced fit via reduced-order kinetics. The present analysis is formulated in the deterministic mass-action regime appropriate for large-copy-number systems; extending EOT to stochastic low-copy-number settings is a natural future direction. In compartments with transport bottlenecks or anomalous diffusion, or in systems with cooperative assembly, effective rates may be time-dependent. Extending the theory to such non-Markovian dynamics— for example, using memory kernels or distributed delays—is feasible and would refine long-time predictions. Target heterogeneity and spatial microdomains can be represented by mixture models, though careful experimental design remains essential.

Looking forward, EOT offers an exposure-aware, mechanism-aware calculus for target engagement. The monotone dependence of EOT_*∞*_ on *k*_3_*/k*_2_ yields an operative design principle: match medicinal-chemistry effort (on *k*_2_) and formulation/PK effort (on *k*_3_) to the regime the system inhabits, rather than optimizing one axis in isolation. Coupling this framework to standard PK/PD models should produce tractable rules for dosage, scheduling, and scaffold selection, with clear, testable predictions across parameter domains.

## Code Availability

The Python scripts used to generate all numerical figures in this article are available in a public repository on GitHub: https://github.com/schnell-lab/ehrlich-occupancy-time. The repository includes a `README` file with installation instructions and step-by-step guidance for reproducing each figure, along with a `requirements.txt` file specifying the exact software versions used.

## Author Contributions Statement

J.E., S.S. and S.W. made substantial contributions to the conception and design of the work, analysis and interpretation of the results. S.S., S.W. and J.E. wrote the manuscript and S.S. and J.E. prepared the figures. All authors reviewed and approved the manuscript.

## Acknowledgement

We thank both anonymous reviewers for thoroughly reading the manuscript, and for their constructive comments and valuable suggestions. Special thanks are due to the reviewer who showed how to streamline some mathematical arguments.

## Appendices A Alternative exponential bounds via variation of constants

In Section 4, we derived bounds on EOT_*∞*_ using comparison with constant-coefficient linear equations. Here we present an alternative approach using variation of constants, which yields bounds with exponential correction factors. These bounds can be sharper in certain parameter regimes.

### A.1 Setup

Recall from Equation (51) that the complex concentration satisfies:

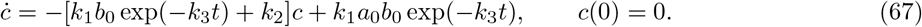

The homogeneous equation 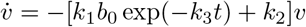 has solution:

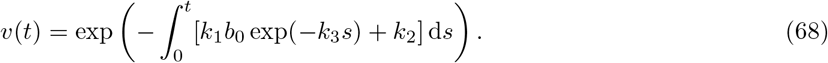

Evaluating the integral:

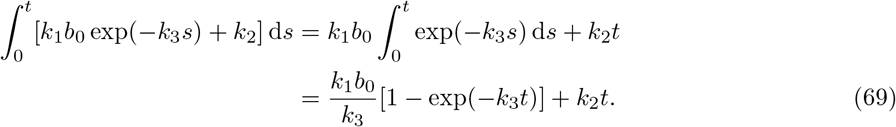

Therefore:

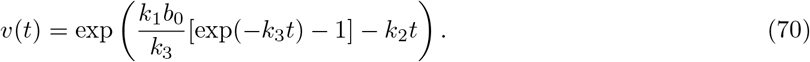

This can be rewritten as:

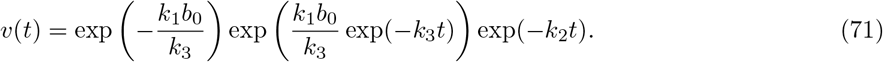

### A.2 Bounds on the Homogeneous Solution

For *t* ≥ 0, we have 0 ≤ exp(−*k*_3_*t*) ≤ 1, which implies:

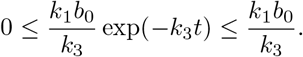

Since the exponential function is monotonically increasing:

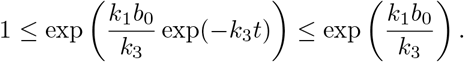

Substituting into (71):

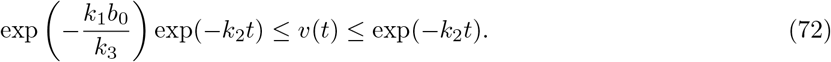

For the inverse:

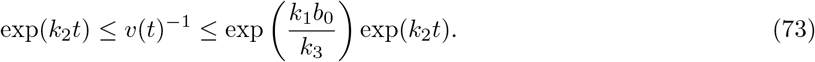

### A.3 Solution via Variation of Constants

The solution to (67) can be written as *c*(*t*) = *w*(*t*)*v*(*t*), where *w*(*t*) satisfies:

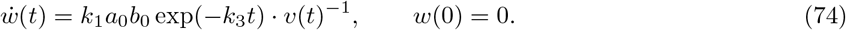

**Upper Bound on** *w*(*t*). Using the upper bound from (73):

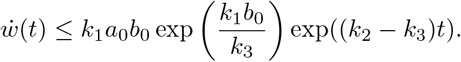

Integrating from 0 to *t* (for *k*_2_

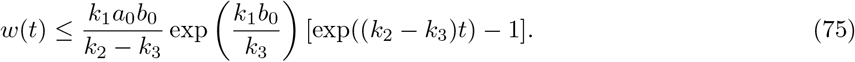

**Lower Bound on** *w*(*t*). Using the lower bound from (73):

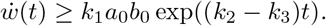

Integrating:

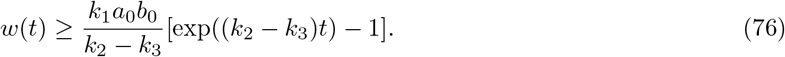

### A.4 Bounds on *c*(*t*)

Combining the bounds on *w*(*t*) and *v*(*t*):

**Upper Bound**. Using (75) and the upper bound on *v*(*t*) from (72):

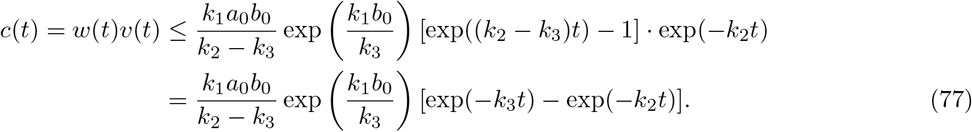

**Lower Bound**. Using (76) and the lower bound on *v*(*t*) from (72):

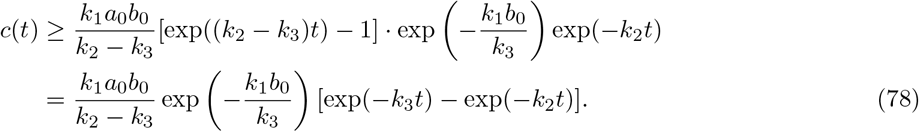

### A.5 Bounds on EOT_*∞*_

Integrating (77) and (78) over [0, ∞) and dividing by *a*_0_ yields the exponential bounds.

#### Upper bound

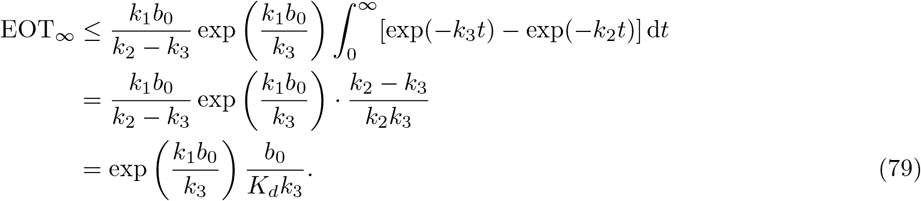

#### Lower bound

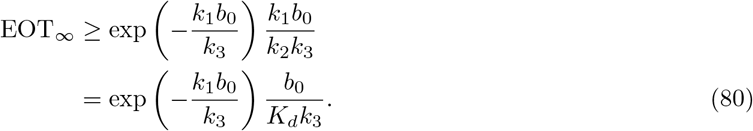

Combining these results:

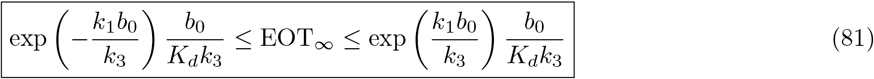

### A.6 Comparison with Standard Bounds

In Section 4, we derived the standard bounds:

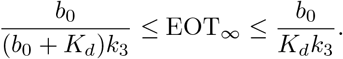

#### Upper bounds

The standard upper bound is *b*_0_*/*(*K*_*d*_*k*_3_), while the exponential upper bound is exp(*k*_1_*b*_0_*/k*_3_)· *b*_0_*/*(*K*_*d*_*k*_3_). Since exp(*k*_1_*b*_0_*/k*_3_) ≥ 1, the **standard upper bound is always tighter**.

#### Lower bounds

The standard lower bound is *b*_0_*/*[(*b*_0_ + *K*_*d*_)*k*_3_], while the exponential lower bound is exp(−*k*_1_*b*_0_*/k*_3_) · *b*_0_*/*(*K*_*d*_*k*_3_). The standard lower bound is tighter when:

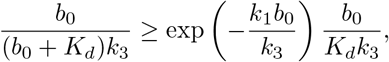

which simplifies to:

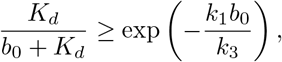

or equivalently:

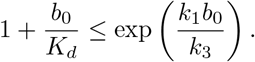

Using the inequality 1+*x* ≤ exp(*x*) for all *x* ≥ 0, this condition is satisfied whenever *b*_0_*/K*_*d*_ ≤ (*b*_0_*/K*_*d*_)·(*k*_2_*/k*_3_), which holds when *k*_2_ ≥ *k*_3_—that is, when dissociation is at least as fast as elimination. Since this is the biologically common regime, the **standard lower bound is typically tighter**.

##### Remark 7

*The exponential bounds may be useful in specialized scenarios where k*_3_ *> k*_2_ *(elimination faster than dissociation) or when asymptotic analysis is required. For most practical applications, the standard bounds from Section 4 are preferred*.

## B Practical application of the EOT framework

Application of the framework requires estimates of *k*_1_, *k*_2_, the relevant initial free-drug concentration *b*_0_, and, for open systems, the effective elimination rate *k*_3_ in the biological compartment of interest. In closed systems, rEOT is approximated by Equation (24); for induced-fit systems by Equation (41) with 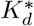 from Equation (40); and for systems with first-order drug removal, EOT_*∞*_ is bounded by Equation (63). When all parameters are known, the full system (43) can also be integrated numerically.

For experimental validation, one measures a time course of fractional occupancy *f*_exp_(*t*) using a targetengagement assay appropriate to the system, and computes

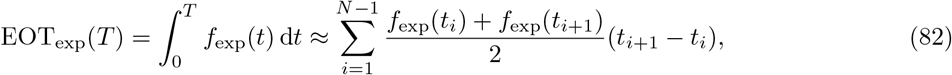

which can be compared directly with the model prediction.

A useful practical diagnostic is the ratio *k*_3_*/k*_2_. When *k*_3_ ≪ *k*_2_, cumulative occupancy is limited mainly by binding kinetics and reducing *k*_2_ can materially increase EOT. When *k*_3_ ≳ *k*_2_, clearance or efflux limits occupancy, so improving exposure may be more effective than further reducing *k*_2_. For induced-fit systems, the same logic applies after replacing *K*_*d*_ by 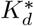.

As a representative open-system calculation, if 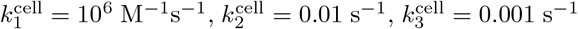, and *b*_0_ = 1 *µ*M, then *K*_*d*_ = 0.01 *µ*M and Equation (63) gives 990 s ≤ EOT_*∞*_ ≤ 100,000 s. Numerical integration of system (43) yields EOT_*∞*_ ≈ 5,160 s, confirming that the analytical bounds correctly bracket the exact value.

Assay-specific protocols, additional illustrative figures, and extended workflow examples have been moved to the **Supplementary Material**.

## Footnotes

1 In the present paper, the right-hand sides are polynomial, derived from mass action kinetics, but the approach applies to more general kinetics as well.

## References

[1] R. A. Copeland, D. L. Pompliano, and T. D. Meek. Drug-target residence time and its implications for lead optimization. Nat. Rev. Drug Discov., 5(9):730–739, 2006. doi: 10.1038/nrd2082.

[2] D. C. Swinney. Biochemical mechanisms of drug action: what does it take for success? Nat. Rev. Drug Discov., 3(9):801–808, 2004. doi: 10.1038/nrd1500.

[3] P. J. Tummino and R. A. Copeland. Residence time of receptor-ligand complexes and its effect on biological function. Biochemistry, 47(20):5481–5492, 2008. doi: 10.1021/bi8002023.

[4] S. Núñez, J. Venhorst, and C. G. Kruse. Target-drug interactions: first principles and their application to drug discovery. Drug Discov. Today, 17(1-2):10–22, 2012. doi: 10.1016/j.drudis.2011.06.013.

[5] P. Ehrlich. Chemotherapeutics: scientific principles, methods and results. Lancet, 182(4694):445–451, 1913.

[6] R. A. Copeland. The drug-target residence time model: a 10-year retrospective. Nat. Rev. Drug Discov., 15(2):87–95, 2016. doi: 10.1038/nrd.2015.18.

[7] R. A. Copeland. Conformational adaptation in drug-target interactions and residence time. Future Med. Chem., 3(12):1491–1501, 2011. doi: 10.4155/fmc.11.112.

[8] J. M. Bradshaw, J. M. McFarland, V. O. Paavilainen, A. Bisconte, D. Tam, V. T. Phan, S. Romanov, D. Finkle, J. Shu, V. Patel, T. Ton, X. Li, D. G. Loughhead, P. A. Nunn, D. E. Karr, M. E. Gerritsen, J. O. Funk, T. D. Owens, E. Verner, K. A. Brameld, R. J. Hill, D. M. Goldstein, and J. Taunton. Prolonged and tunable residence time using reversible covalent kinase inhibitors. Nat. Chem. Biol., 11 (7):525–531, 2015. doi: 10.1038/nchembio.1817.

[9] G. K. Walkup, Z. You, P. L. Ross, E. K. H. Allen, F. Daryaee, M. R. Hale, J. O’Donnell, D. E. Ehmann, V. J. A. Schuck, E. T. Buurman, A. L. Choy, L. Hajec, K. Murphy-Benenato, V. Marone, S. A. Patey, L. A. Grosser, M. Johnstone, S. G. Walker, P. J. Tonge, and S. L. Fisher. Translating slow-binding inhibition kinetics into cellular and in vivo effects. Nat. Chem. Biol., 11(6):416–423, 2015. doi: 10.1038/nchembio.1796.

[10] S. Davoodi, F. Daryaee, A. Chang, S. G. Walker, and P. J. Tonge. Correlating drug-target residence time and post-antibiotic effect: Insight into target vulnerability. ACS Infect. Dis., 6(4):629–636, 2020. doi: 10.1021/acsinfecdis.9b00484.

[11] G. Vauquelin. Effects of target binding kinetics on in vivo drug efficacy: koff, kon and rebinding. Br. J. Pharmacol., 173(15):2319–2334, 2016. doi: 10.1111/bph.13504.

[12] R. H. A. Folmer. Drug target residence time: a misleading concept. Drug Discov. Today, 23(1):12–16, 2018. doi: 10.1016/j.drudis.2017.07.016.

[13] T. M. Apostol. Mathematical Analysis. Addison-Wesley, Reading, MA, 2nd edition, 1974.

[14] W. Rudin. Real and Complex Analysis. McGraw-Hill Book Co., New York, 3rd edition, 1987.

[15] R. A. Copeland. Evolution of the drug-target residence time model. Expert Opin. Drug Discov., 16(12): 1441–1451, 2021. doi: 10.1080/17460441.2021.1948997.

[16] P. Csermely, R. Palotai, and R. Nussinov. Induced fit, conformational selection and independent dynamic segments: an extended view of binding events. Trends Biochem. Sci., 35(10):539–546, 2010. doi: 10.1016/j.tibs.2010.04.009.

[17] A. D. Vogt and E. Di Cera. Conformational selection or induced fit? A critical appraisal of the kinetic mechanism. Biochemistry, 53(36):5723–5729, 2014. doi: 10.1021/bi500913n.

[18] W. Walter. Ordinary Differential Equations. Springer-Verlag, New York, 1998. doi: 10.1007/978-1-4612-0601-9.

[19] Pauli Virtanen, Ralf Gommers, Travis E. Oliphant, Matt Haberland, Tyler Reddy, David Cournapeau, Evgeni Burovski, Pearu Peterson, Warren Weckesser, Jonathan Bright, Stéfan J. van der Walt, Matthew Brett, Joshua Wilson, K. Jarrod Millman, Nikolay Mayorov, Andrew R. J. Nelson, Eric Jones, Robert Kern, Eric Larson, C. J. Carey, İlhan Polat, Yu Feng, Eric W. Moore, Jake VanderPlas, Denis Laxalde, Josef Perktold, Robert Cimrman, Ian Henriksen, E. A. Quintero, Charles R. Harris, Anne M. Archibald, Antônio H. Ribeiro, Fabian Pedregosa, Paul van Mulbregt, and SciPy 1.0 Contributors. SciPy 1.0:SciPy 1.0: Fundamental algorithms for scientific computing in Python. Nature Methods, 17:261–272, 2020. doi: 10.1038/s41592-019-0686-2.

