## Supplementary Material for "Ehrlich occupancy time: Beyond *k*_off_ to a complete residence time framework"


This supplementary document collects the application-oriented material moved out of Appendix B of the main manuscript in order to keep the article focused on the mathematical framework. The compact computational workflow, the experimental integration formula for $\text{EOT}_{\text{exp}}$, and the representative open-system calculation are retained in the main paper. The present supplement gathers assay-oriented guidance for estimating $k_1$, $k_2$, and $k_3$, two additional illustrative figures, experimental validation modalities, and an extended optimization workflow.

Throughout, we use the main-text expressions

$$\text{rEOT} \approx \frac{b_0}{K_{\text{d}} + b_0}, \qquad \frac{b_0}{(b_0 + K_{\text{d}})k_3} \leq \text{EOT}_\infty \leq \frac{b_0}{K_{\text{d}}\, k_3}, \qquad K_d^* = \frac{K_{\text{d}}}{1 + k_3/k_4},$$

for closed systems under pseudo-first-order conditions, open systems with first-order drug removal, and induced-fit systems, respectively.

### S1 Parameter estimation

Applying the EOT framework requires estimates of the association rate constant $k_1$, the dissociation rate constant $k_2$, and, for open systems, the effective elimination rate constant $k_3$ in the biological compartment of interest. The protocols below are intended as practical routes to obtaining these quantities in biochemical, cellular, and in vivo settings.

#### S1.1 Measuring $k_1$ and $k_2$ in biochemical systems

**Surface plasmon resonance (SPR).** Purified target protein is immobilized on a sensor chip, and drug solutions are flowed across the surface at 5–7 concentrations spanning roughly $0.1K_\mathrm{d}$ to $10K_\mathrm{d}$. Association and dissociation phases are recorded and fit globally to a 1:1 Langmuir model,

$$R(t) = R_\mathrm{max}\big[1 - \exp(-(k_1 C + k_2)t)\big] \quad \text{(association)}, \qquad R(t) = R_0 \exp(-k_2 t) \quad \text{(dissociation)}.$$

SPR is often the most direct route to separate estimates of $k_1$ and $k_2$, particularly when the system is close to ideal 1:1 binding [1].

**Bio-layer interferometry (BLI).** BLI uses fiber-optic sensors rather than microfluidic flow cells. It can be operationally simpler than SPR and is often useful for rapid triage or for work with crude lysates. Its lower sensitivity for extremely tight binders, however, can make it less suitable when $K_\mathrm{d}$ is well below 1 nM.

**Stopped-flow fluorescence.** When the relevant kinetic phases are too fast for reliable SPR resolution, stopped-flow fluorescence provides an alternative. This is especially useful when either the ligand or the target can be monitored spectroscopically and the dominant kinetic phase is on the millisecond timescale.

**Quality control.** The kinetically inferred dissociation constant $K_\mathrm{d}^\mathrm{kinetic} = k_2/k_1$ should be checked against an independently measured equilibrium estimate. Discrepancies larger than about 2–3-fold are often a sign of non-1:1 stoichiometry, mass-transport artifacts, heterogeneous target preparation, or unresolved multiphasic binding.

#### S1.2 Measuring cellular $k_1^\mathrm{cell}$ and $k_2^\mathrm{cell}$

Cellular rate constants can differ materially from biochemical values because the effective free drug concentration, diffusion environment, compartmentalization, and competition with endogenous ligands differ from purified in vitro assays [2].

**Washout experiments.** Cells are exposed to drug until a quasi-steady occupancy level is reached, then extracellular drug is removed by rapid media exchange. The ensuing decay in intracellular target engagement can be monitored with fluorescent ligands, NanoBRET target-engagement assays, or CETSA. Fitting the post-washout decay provides an operational estimate of $k_2^\mathrm{cell}$.

**Pulse-chase competition.** After equilibration with a labeled ligand, a saturating concentration of unlabeled competitor is added. If competitor binding is sufficiently fast and the competitor concentration greatly exceeds $K_\mathrm{d}$, the observed recovery is dominated by dissociation of the original ligand and again provides an estimate of $k_2^\mathrm{cell}$.

**Association measurements in cells.** Cellular estimates of $k_1^\mathrm{cell}$ are typically less direct than cellular estimates of $k_2^\mathrm{cell}$. A practical route is to record the occupancy time course at several drug concentrations and fit the resulting family of trajectories jointly. When an independent cellular $K_\mathrm{d}^\mathrm{cell}$ estimate is available, one may also use the relation $k_1^\mathrm{cell} = k_2^\mathrm{cell}/K_\mathrm{d}^\mathrm{cell}$ as an approximation.

#### S1.3 Measuring $k_3$

**Liver microsome stability.** A standard first screen is to incubate drug (for example at 1 $\mu$M) with pooled liver microsomes and quantify parent compound over time by LC–MS/MS. Fitting a first-order decay gives an effective microsomal removal constant. This assay is convenient, but it captures only part of whole-organism elimination.

**Hepatocyte stability.** Cryopreserved hepatocytes often provide a more physiological estimate because they preserve transporter activity and intracellular cofactors. Variability is usually higher than with microsomes, but the readout is often closer to the integrated effect of metabolism and cellular transport.

**In vivo pharmacokinetics.** For a systemic estimate, concentration–time profiles after an intravenous dose can be fit by one- or multi-compartment pharmacokinetic models. In the one-compartment limit, $k_3$ corresponds to clearance divided by volume of distribution.

**Intracellular $k_3$.** For intracellular targets, the relevant elimination constant is frequently not the plasma value but the effective intracellular decay rate. This quantity can reflect active efflux, passive diffusion back across the membrane, intracellular metabolism, or trapping in subcellular compartments. Direct measurement by cell lysis plus LC–MS, or by calibrated live-cell fluorescence methods, is often more informative than relying on plasma kinetics alone.

**Illustrative example.** Some kinase inhibitors show plasma half-lives of order 10 h but intracellular half-lives closer to 2 h because of transporter-mediated efflux. In such cases, intracellular $k_3$ rather than plasma $k_3$ is the limiting quantity for occupancy-based interpretation.

Table S1: Typical order-of-magnitude ranges for kinetic and elimination parameters encountered in drug-discovery settings.

| **Parameter** | **Typical range** |
|---|---|
| $k_1$ | $10^4$ to $10^7$ $M^{-1}s^{-1}$ (diffusion-limited upper bound $\sim 10^9$) |
| $k_2$ | $10^{-4}$ to 1 $s^{-1}$ (ultra-tight to weak binders) |
| $k_3$ | $10^{-5}$ to 1 $s^{-1}$ (persistent to rapidly cleared drug) |

### S2 Additional experimental implications and illustrative figures

The main manuscript derives the governing expressions and provides the principal numerical validation. The present section records several additional qualitative implications that may be useful when designing experiments around the theory.

#### S2.1 Concentration dependence in open systems

The open-system bounds imply a crossover in cumulative occupancy as drug concentration increases. When $b_0 \ll K_\mathrm{d}$, the lower bound behaves as

$$\frac{b_0}{(b_0 + K_\mathrm{d})k_3} \approx \frac{k_1 b_0}{k_2 k_3},$$

so cumulative occupancy grows approximately linearly with concentration. When $b_0 \gg K_\mathrm{d}$, the same lower bound approaches $1/k_3$, so the concentration dependence saturates.

This crossover is already visible in the lower analytical bound plotted in Figure S1. The exact $\mathrm{EOT}_\infty$ lies above the lower-bound curves, but exhibits the same qualitative transition from a concentration-sensitive regime to a clearance-limited plateau.

**Illustrative protocol.** A practical experiment is to select a well-characterized drug–target pair, measure occupancy time courses over a concentration range spanning roughly $0.1K_\mathrm{d}$ to $10K_\mathrm{d}$, and compute $\mathrm{EOT}(T)$ over a fixed observation window. The expected behavior is a low-concentration regime where increasing dose materially increases occupancy time, followed by a high-concentration regime where further increases in $b_0$ yield diminishing gains because elimination becomes the dominant limitation.

#### S2.2 Dependence on the elimination rate

For fixed $k_1$ and $k_2$, both analytical bounds scale inversely with $k_3$. A reduction of the relevant elimination constant by a factor of 10 therefore increases the predicted occupancy-time scale by the same factor. Comparisons across species, transporter knockouts, or the presence versus absence of metabolic inhibitors are natural experimental settings in which to test this dependence.

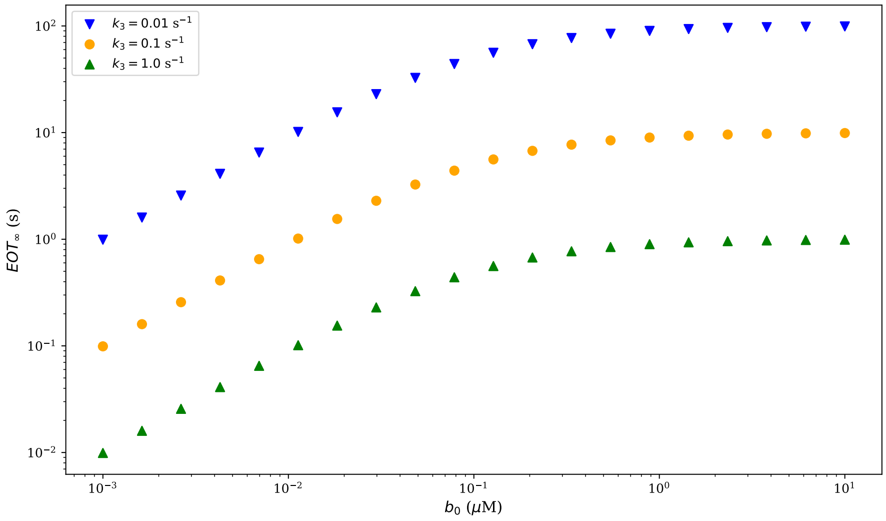


Figure S1: Illustrative concentration dependence of the lower analytical bound on $\mathrm{EOT}_\infty$. The plotted quantity is $b_0/[(b_0 + K_\mathrm{d})k_3]$ for three elimination rates. At low concentration ($b_0 \ll K_\mathrm{d}$), the bound scales approximately linearly with $b_0$; at high concentration ($b_0 \gg K_\mathrm{d}$), it approaches the clearance-limited plateau $1/k_3$. Parameters: $k_1 = 10^6\ \mathrm{M}^{-1}\mathrm{s}^{-1}$, $k_2 = 0.1\ \mathrm{s}^{-1}$, and hence $K_\mathrm{d} = 0.1\ \mu\mathrm{M}$.

##### S2.3 Induced fit

For induced-fit systems, the equilibrium occupancy is controlled by the effective dissociation constant $K_d^* = K_\mathrm{d}/(1+k_3/k_4)$. Increasing the ratio $k_3/k_4$ therefore shifts occupancy upward even when the initial encounter complex has the same intrinsic $K_\mathrm{d}$.

**Illustrative protocol.** For a target with a known induced-fit mechanism, one may first estimate the initial binding constant from rapid equilibrium measurements, then estimate the isomerization rates from stopped-flow, temperature-jump, or multiphasic dissociation experiments. The occupancy predicted from $K_d^*$ can then be compared with direct cellular target-engagement measurements.

##### S2.4 Bound accuracy

A direct way to assess the analytical bounds is to combine measured $k_1$, $k_2$, $k_3$, and $b_0$ values with an experimentally reconstructed occupancy curve. The resulting empirical EOT should fall between the lower and upper bounds if the pseudo-first-order approximation is adequate and the first-order removal model is appropriate. Extensive parameter sweeps and exact numerical calculations are provided in Figures 2 and 3 of the main manuscript.

#### S3 Experimental validation modalities

Once the relevant kinetic parameters have been estimated, the main practical question is how to measure or infer the occupancy curve $f(t)$ in the biological system of interest. The main manuscript gives the numerical integration formula for converting such measurements into $\mathrm{EOT}_\mathrm{exp}$; the paragraphs below summarize common assay modalities.

##### S3.1 Direct target-occupancy measurements

**CETSA.** Drug binding can stabilize a protein against thermal denaturation. Cells are treated with compound, optionally washed, then exposed to a temperature gradient. The residual soluble target is quantified

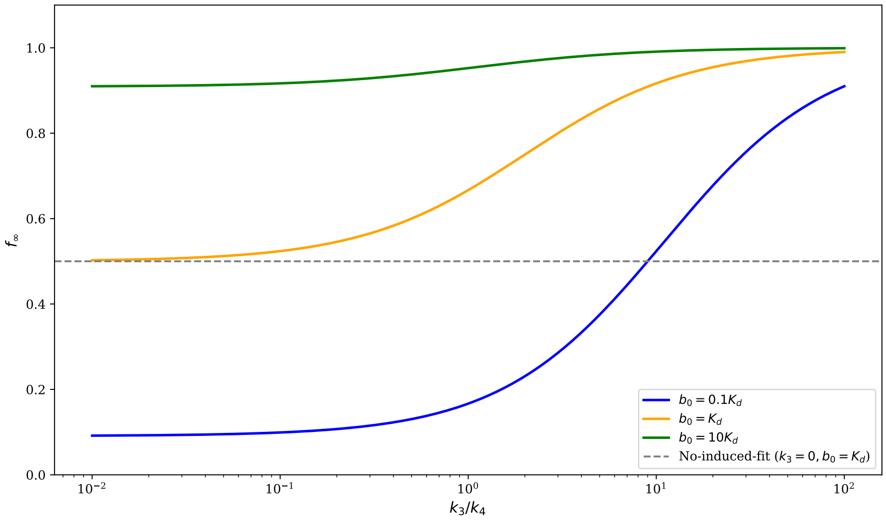


Figure S2: Illustrative effect of induced fit on the equilibrium occupancy fraction $f_\infty$. As the isomerization ratio $k_3/k_4$ increases, the effective dissociation constant $K_d^*$ decreases and occupancy rises. The effect is modest when $b_0 \ll K_\mathrm{d}$ or $b_0 \gg K_\mathrm{d}$, and is most visually pronounced for intermediate concentration. Parameters: $K_\mathrm{d} = 1\ \mu$M.

after lysis. A time course of thermal stabilization after washout provides an operational readout of how long the target remains occupied.

**NanoBRET target engagement.** A NanoLuc-tagged target and a fluorescent tracer provide a live-cell, competition-based readout of target engagement. The method is especially attractive when repeated measurements in living cells are needed, because association, dissociation, and washout recovery can all be followed in the same basic experimental framework.

**Activity-based proteomics.** For enzymes such as kinases or proteases, a post-washout labeling step with an active-site-directed probe can report residual target accessibility. Lower probe labeling implies greater persistence of occupancy by the test compound.

#### S3.2 Indirect pharmacodynamic readouts

When direct occupancy measurements are impractical, downstream biomarkers can provide useful surrogates, with the usual caution that signaling feedback and network dynamics can decouple biomarker recovery from target disengagement.

**Kinase inhibitors.** A common strategy is to monitor recovery of substrate phosphorylation after washout. The approach is most informative when the biomarker is tightly linked to the inhibited kinase and feedback is limited.

**GPCR antagonists.** Cells or tissues are pre-treated with antagonist, washed, and then re-stimulated with agonist at later times. Recovery of the agonist response provides an indirect estimate of antagonist dissociation and loss of occupancy.

**Antibiotics.** The post-antibiotic effect (PAE) provides an indirect duration metric for antibacterial agents. In systems where the target vulnerability is well understood, PAE duration can be interpreted alongside residence-time and occupancy-time ideas [3].

##### S3.3 Assessing agreement between prediction and experiment

As a practical rule of thumb, agreement within about 2-fold between predicted and measured occupancy time is usually strong support for the working model. Discrepancies in the 2–5-fold range often indicate uncertainty in $k_3$, assay-to-assay variation, or moderate model mismatch. Larger discrepancies suggest additional mechanisms such as target turnover, multiple binding modes, active transport, or active metabolites.

#### S4 Optimization considerations

The main manuscript argues that occupancy-aware optimization depends on the relative magnitudes of $k_2$ and $k_3$. The paragraphs below translate that idea into medicinal-chemistry and assay-planning language.

##### S4.1 When binding kinetics are limiting

If $k_3 \ll k_2$, drug remains available long enough that slowing dissociation can materially increase cumulative occupancy. In that regime, medicinal-chemistry strategies aimed at reducing $k_2$—for example by increasing buried surface area, improving polar complementarity in a deep pocket, or exploiting conformational trapping—are likely to be worthwhile.

##### S4.2 When pharmacokinetics are limiting

If $k_3 \gg k_2$, further decreases in $k_2$ deliver diminishing returns because drug is lost from the relevant compartment before the advantage of slower dissociation can be fully realized. In this regime, the main opportunities lie in reducing effective elimination: improving metabolic stability, reducing efflux, or altering distribution.

##### S4.3 Formulation strategies

When the binding scaffold is already satisfactory but exposure is short-lived, formulation changes that effectively reduce the relevant $k_3$ can increase occupancy time approximately in proportion to the gain in persistence. Sustained-release formulations, depot delivery, nanoparticle encapsulation, and related strategies are therefore natural complements to chemistry optimization.

#### S5 Extended illustrative workflow

The following hypothetical kinase-inhibitor program is included as an extended illustration of how the EOT framework can be used operationally.

1. A screening hit shows biochemical $K_\mathrm{d} = 50$ nM but cellular $\mathrm{IC}_{50} = 500$ nM.
2. Kinetic characterization gives $k_1 = 10^7$ $\mathrm{M}^{-1}\mathrm{s}^{-1}$ and $k_2 = 0.5$ $\mathrm{s}^{-1}$.
3. Washout experiments indicate an effective intracellular elimination constant $k_3^{\mathrm{cell}} = 0.05$ $\mathrm{s}^{-1}$.
4. At $b_0 = 500$ nM, the open-system upper bound gives

$$\mathrm{EOT}^u_\infty = \frac{500\,\mathrm{nM}}{50\,\mathrm{nM} \times 0.05\,\mathrm{s}^{-1}} = 200\,\mathrm{s}.$$

5. The interpretation is that rapid intracellular loss, rather than poor affinity, is the main limitation on cumulative occupancy.
6. Additional profiling identifies a high efflux ratio, suggesting transporter-mediated loss from the cell.
7. A next chemistry round reduces basicity to lessen transporter recognition.
8. The follow-up compound yields $k_3^{\mathrm{cell}} = 0.005$ $\mathrm{s}^{-1}$, a 10-fold improvement in persistence.

9. The corresponding upper-bound estimate increases 10-fold to 2000 s.

10. Follow-up target-engagement experiments are then used to test whether the longer intracellular persistence translates into longer occupancy and improved cellular potency.

This workflow is intentionally schematic, but it illustrates how the framework helps separate binding-limited problems from exposure-limited problems and therefore guides which optimization lever is most likely to matter.

### Code Availability

The Python scripts used to generate all numerical figures in this article are available in a public repository on GitHub: `https://github.com/schnell-lab/ehrlich-occupancy-time`. The repository includes a `README` file with installation instructions and step-by-step guidance for reproducing each figure, along with a `requirements.txt` file specifying the exact software versions used.

### References


[1] R. A. Copeland, D. L. Pompliano, and T. D. Meek. Drug-target residence time and its implications for lead optimization. *Nat. Rev. Drug Discov.*, 5(9):730–739, 2006. doi: 10.1038/nrd2082. Erratum in: Nat. Rev. Drug Discov. 2007, 6(3), 249.

[2] G. Vauquelin. Effects of target binding kinetics on in vivo drug efficacy: $k_{\text{off}}$, $k_{\text{on}}$ and rebinding. *Br. J. Pharmacol.*, 173(15):2319–2334, 2016. doi: 10.1111/bph.13504.

[3] S. Davoodi, F. Daryaee, A. Chang, S. G. Walker, and P. J. Tonge. Correlating drug-target residence time and post-antibiotic effect: Insight into target vulnerability. *ACS Infect. Dis.*, 6(4):629–636, 2020. doi: 10.1021/acsinfecdis.9b00484.
